# Model-free Estimation of Tuning Curves and Their Attentional Modulation, Based on Sparse and Noisy Data

**DOI:** 10.1101/026682

**Authors:** Markus Helmer, Vladislav Kozyrev, Valeska Stephan, Stefan Treue, Theo Geisel, Demian Battaglia

**Affiliations:** Max Planck Institute for Dynamics and Self-Organization, Department of Nonlinear Dynamics, Göttingen, Germany; Bernstein Center for Computational Neuroscience, Göttingen, Germany; Institute of Neuroinformatics, Ruhr-University Bochum, Germany; Cognitive Neuroscience Laboratory, German Primate Center, Göttingen, Germany; Institut de Neurosciences des Systèmes, Aix-Marseille Université, Marseille, France

## Abstract

Tuning curves are the functions that relate the responses of sensory neurons to various values within one continuous stimulus dimension (such as the orientation of a bar in the visual domain or the frequency of a tone in the auditory domain). They are commonly determined by fitting a model e.g. a Gaussian or other bell-shaped curves to the measured responses to a small subset of discrete stimuli in the relevant dimension. However, as neuronal responses are irregular and experimental measurements noisy, it is often difficult to determine reliably the appropriate model from the data. We illustrate this general problem by fitting diverse models to representative recordings from area MT in rhesus monkey visual cortex during multiple attentional tasks involving complex composite stimuli. We find that all models can be well-fitted, that the best model generally varies between neurons and that statistical comparisons between neuronal responses across different experimental conditions are affected quantitatively and qualitatively by specific model choices. As a robust alternative to an often arbitrary model selection, we introduce a model-free approach, in which features of interest are extracted directly from the measured response data without the need of fitting any model. In our attentional datasets, we demonstrate that data-driven methods provide descriptions of tuning curve features such as preferred stimulus direction or attentional gain modulations which are in agreement with fit-based approaches when a good fit exists. Furthermore, these methods naturally extend to the frequent cases of uncertain model selection. We show that model-free approaches can identify attentional modulation patterns, such as general alterations of the irregular shape of tuning curves, which cannot be captured by fitting stereotyped conventional models. Finally, by comparing datasets across different conditions, we demonstrate effects of attention that are cell-and even stimulus-specific. Based on these proofs-of-concept, we conclude that our data-driven methods can reliably extract relevant tuning information from neuronal recordings, including cells whose seemingly haphazard response curves defy conventional fitting approaches.

## Introduction

Tuning curves represent a sensory neuron’s response profile to a continuous stimuli parameter (such as orientation, direction of motion, or spatial frequency in the visual domain) and are an ubiquitous tool in neuroscience. In order to describe such selectivities in simple terms, tuning curves are commonly modeled by fitting suitable shape functions to the data, such as, for example, Gaussian distributions [1–4], arbitrary polynomials [5] or generic Fourier series [6] for orientation and direction tuning curves, or other smooth functions like splines [7] for non bell-shaped tuning profiles.

However, the measured tuning curve shapes are tremendously variable and they often deviate from the assumed reference shape [8–11]. More than thirty years ago, De Valois, Yund and Hepler already observed that no single function could adequately describe the tuned response of all cells [8] they had recorded. A few years later, Swindale found that several models fit his data equally well and further questioned the existence of a single all-encompassing model, observing, moreover, that the preferred stimulus deduced from a fitted tuning curve depended on the chosen model [9]. These and other studies thus manifest that the choice of a model to fit can affect the conclusions reached about tuning properties and their contextual modulations.

Here, we will first systematically investigate the problem of ambiguous model selection, highlighting the possible consequences of the choice of a “wrong” model. We will then introduce alternative methods which allow the extraction of features of interest directly from the measured neuronal responses, without the need of fitting any model to the empirical neuronal responses to a small subset of possible values from the continuous stimulus dimension. To illustrate the applicability and the heuristic power of our data-driven methods, we will carry out analyses of attentional modulation effects in single unit recordings in the middle temporal visual area (MT) of four rhesus monkeys, where neurons exhibit characteristic direction selective responses to moving visual stimuli [12, 13]. We will consider responses to stimuli consisting of either one or two random dot patterns (RDPs) in the receptive field where the two RDPs could be either spatially separated or transparently superimposed. In these experimental paradigms, tuning curves are expected to display either one or two peaks. The expected effects of attention include gain modulations leading to changes in the amplitude of these response peaks. However, due to the high number of experimental conditions and the difficulty of the animal’s behavioral task, only relatively few trials could be recorded for each stimulus. This limited sampling, combined with the heterogeneity of response profiles, make the measured tuning curves very “noisy”. The dataset, thus, besides its intrinsic interest, provides a perfect test-bed to reveal the drawbacks of fitting techniques and to benchmark alternative methods. We will discuss two complementary strategies in detail.

First, we will parse the trial-averaged responses of single neurons to obtain, through a set of algorithmic rules, a list of features characterizing the neuron’s response profile. For instance, a set of rules will be used to estimate the direction of a cell’s preferred stimulus, solely based on the data points themselves. Analogously, other rules will be used to capture into suitable index quantities generic variations of the average response profile of a cell across different experimental conditions (e.g. different types of stimuli, or targets of attention). As a shared prerequisite, all these rules must be able to operate by just receiving as input the set of average responses to each of the different stimuli from the small set used in the experiment. Although such an approach might seem too coarse compared to continuous interpolations, we will show an excellent agreement between the conclusions reached by feature extraction and fitting methods, whenever reliable model-based estimates can be derived. In addition, the same feature extraction rules will straightforwardly generalize even to the most irregular tuning curves, for which model fitting would be questionable. By the same token, the unbounded flexibility in rule design will allow the extraction of ad hoc features, revealing aspects of shape and shape change which elude a parameterization in terms of conventional fitted models.

Second, we will make use of the full information conveyed by the stochasticity of individual trials. For each neuron and for each different stimulus direction we will quantify the distribution of responses across trials, and compare them across different experimental conditions. This approach will show that attention frequently significantly modulates the response of a cell only for specific stimulus directions rather than a general modulation across the whole tuning profile.

Thus, through proof-of-concept analyses, we demonstrate the potential of data-driven methods for harnessing the heterogeneity of tuned responses. We prove that our approach robustly captures complex effects of attention, avoiding excessive reliance on illustrative well-behaved cases and circumventing the narrow constraints exerted by a fitting framework.

## Results

### The experiment: attentional influences on single-cell responses to composite stimuli

To illustrate drawbacks with fitting approaches and test novel methodology on a concrete representative example, we focused on extra-cellular recordings of single neurons from area MT in the visual cortex of rhesus monkeys [14, 15]. The aim of these recordings was to investigate how attention affects the tuning curves obtained by the simultaneous presentation of two directions of motion within the receptive field of a given neuron.

Neuronal responses were recorded under different conditions (Fig. 1A–E; see Methods for details). In the first experimental paradigm (Fig. 1A–B), random dot patterns (RDPs) moving within two *spatially separated* stationary apertures were used. They were sized and positioned for each recorded neuron to fit within its classical receptive field (RF). In a second experimental paradigm (Fig. 1C–D), both RDPs were fully overlapping. This single aperture contained two sets of random dots, moving transparently in two directions of motion and covered most of the RF. In all conditions a fixed angular separation of 120° between the two RDPs was used.

**Figure 1.**
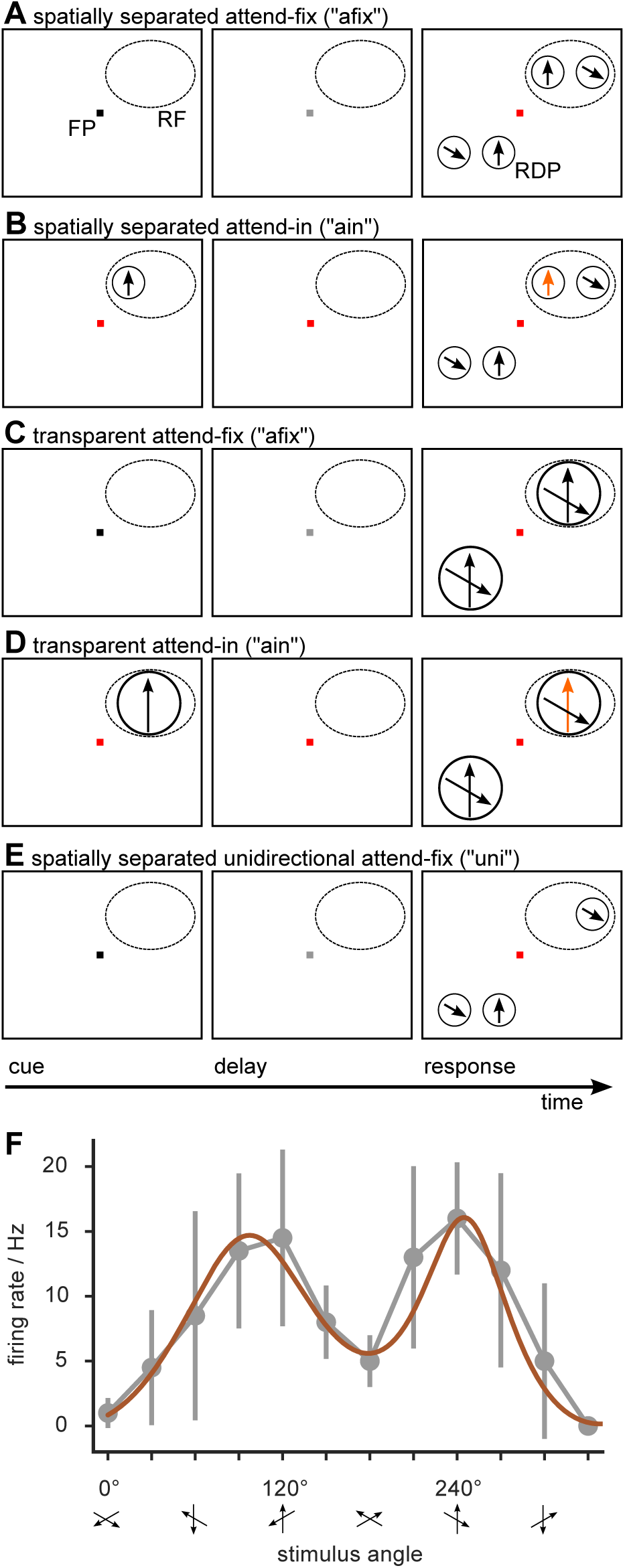
*(following page)*. Attentional experiments. Direction selective responses of MT cells were measured using different direction combinations of stimuli and different attentional conditions. The stimuli in the receptive field (RF) of the recorded cell were either two random dot patterns (RDP) moving in directions 120^*◦*^ apart and placed in spatially separated (panels A–B) or overlapping (panels C–D) apertures or just one single (unidirectional) RDP (panel E). A cue instructed the monkey to attend to either: a luminance change of the fixation point (FP), in the attend-fix condition (afix, panels A and C) and single stimulus (uni, panel E) conditions; or to changes of the direction or velocity of the cued RDP (orange) in the RF, in the attend-in conditions (ain, panels B and D). The transparent uni condition was taken to be the cue-period of the ain condition (panel D). F: Example of a “well-behaved” tuning curve from the spatially separated paradigm in the afix condition. Gray circles denote trial-averaged firing rates and error bars their standard deviation. A sum-of-two-gaussians fit is also shown (brown). The stimulus directions are aligned for each cell, so that the attended direction corresponds to the preferred direction in the uni condition at 240^*◦*^ (see Materials and Methods for details).

In each of the two paradigms, attention could be directed either to a fixation spot outside the recorded RF (condition *“afix”*), or to one of the RDPs inside the RF (condition *“ain”*). In addition, the study included conditions in which only one of the two RDPs was shown while attention was directed to the fixation spot (condition *“uni”*; see Fig. 1E for an explanatory cartoon of a spatially separated *“uni”* trial). Overall, data from six different experimental configurations (spatially separated afix, ain and uni; transparent afix, ain and uni) were analyzed (see also Materials and Methods).

Based on previous studies the *“uni”* experiments should generate unimodal tuning curves, while 120° separation between the spatially separated and transparent *“afix”* and *“ain”* stimuli should result in bimodal tuning curves [16], with the peaks occurring whenever one of the two stimulus components moved in the cell’s preferred direction. Fig. 1F shows an example tuning curve from the spatially separated afix condition. Responses were sampled using 12 different directions across the full 360° range with an angular resolution of 30°. This sampling resolution is typical, as direction-tuning curves are most frequently assessed using 8 or 12 evenly spaced directions to account for the constraints of a behavioral paradigm where the number of trials that can be run is limited. Note that for plotting, stimulus directions were aligned for each cell such that the attended directino in the 240° stimulus was moving in the cell’s preferred direction.

### “Noisy” tuning curves: not one model to rule them all

To model bell-shaped tuning curves, Gaussian curves are commonly used [1–4] and they are usually wrapped, due to the circular nature of the fitted data [17]. This is exemplified in Fig. 1F where we fitted a sum of two Gaussians (brown curve) to a bimodal dataset. In this case, the fit looks adapted to the data, at least according to visual inspection. However, not all the cases are equally “well-behaved”, as evinced, e.g., by comparing the tuning curve of Fig. 1F with a second example in Fig. 2A.

**Figure 2.**
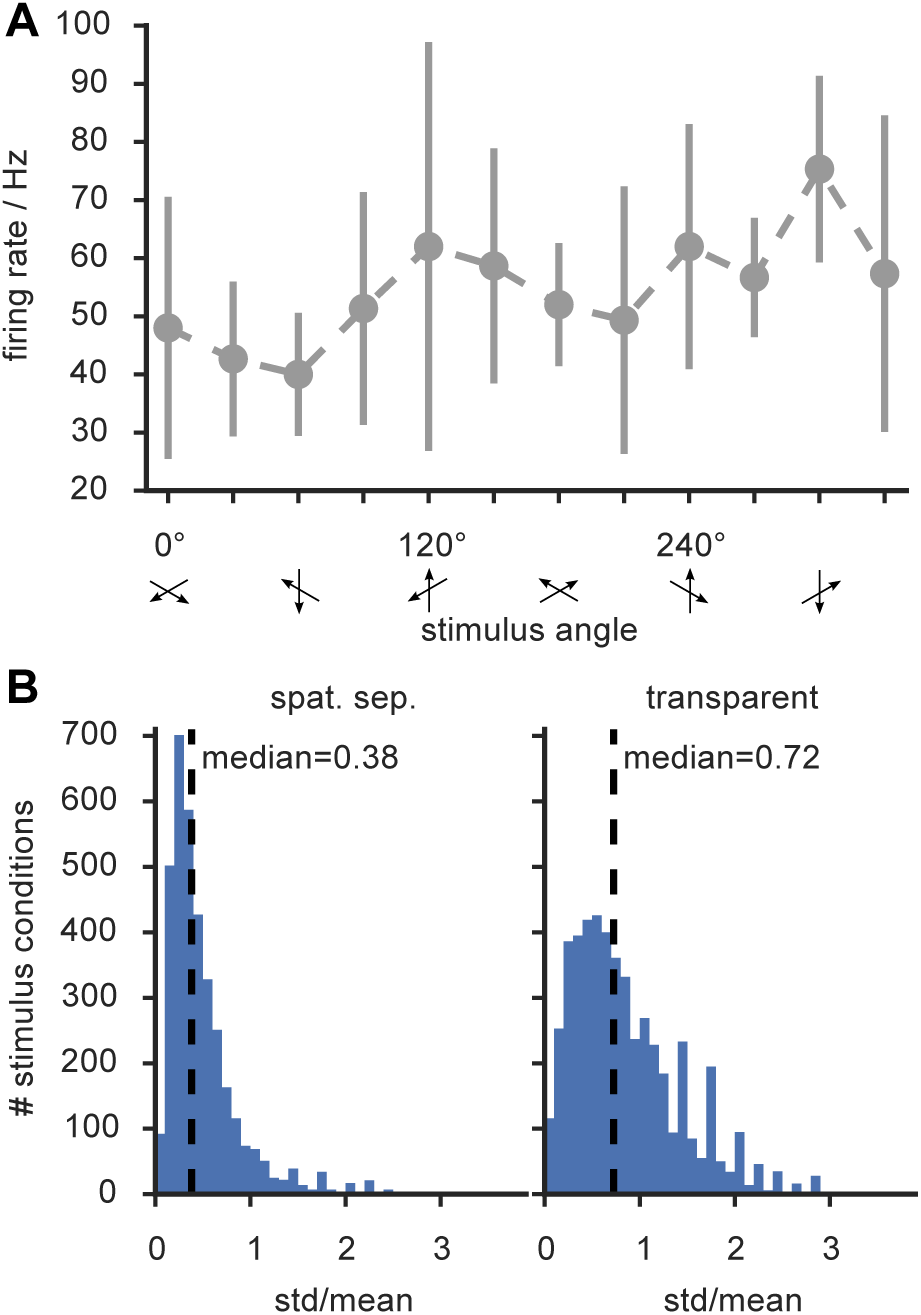
Many tuning curves are not “well-behaved”. A: typical example of tuning curve from the spatially-separated afix condition (compare with Fig. 1F). The shape of the curve—including the position of the two peaks that should be elicited by the composite RDP stimulus—cannot reliably be inferred due to large error bars (std.). B: Histogram of estimated firing rate standard deviations (expressed in relative units, as ratios between std. and a matching mean), obtained by lumping together all stimulus directions and attentional conditions, for the spatially separated (left) and the transparent (right) paradigms. Both these histograms are strongly right-skewed, denoting the existence of cells with highly variable responses to certain stimuli.

Two aspects need to be emphasized. First, the coarse sampling of the response profile because of the 30^*◦*^ separation between measurement points and second, the large trial-by-trial response variability to a given stimulus, visualized by the error bars.

Indeed, the estimated coefficient of variation (i.e. the standard deviation in units of the mean) has a median of 38 % for the spatially separated paradigm and of 72 % for the transparent paradigm, as shown by Fig. 2B. Given this large uncertainty in the data it is thus not surprising that a “well fitted” linear combination of Gaussians lies within the error bars. However, within these large ranges of uncertainty, fitted curves obtained from model functions other than Gaussians could be accommodated as well, and there is no general *a priori* argument that a Gaussian model is the best suited model for such tuning curves.

To corroborate our intuition, we fitted eight different model functions to our data, testing for their compatibility with Gaussian and non-Gaussian shapes (Fig. 3). The used functions were a wrapped Gaussian (brown color) [9], a wrapped Cauchy function (yellow) [17], a symmetric Beta function (violet) [18], a wrapped generalized bell-shaped membership function (pink) [19], a von Mises function [9, 17] (orange), and Fourier series of order 2 (red), 3 (blue) and 4 (green) [6]. Details on the used models are provided in the Materials and Methods section. In Fig. 3A, we show an example neuron for which all tested models provided reasonably looking fits at visual inspection (in the transparent attend fix condition).

**Figure 3.**
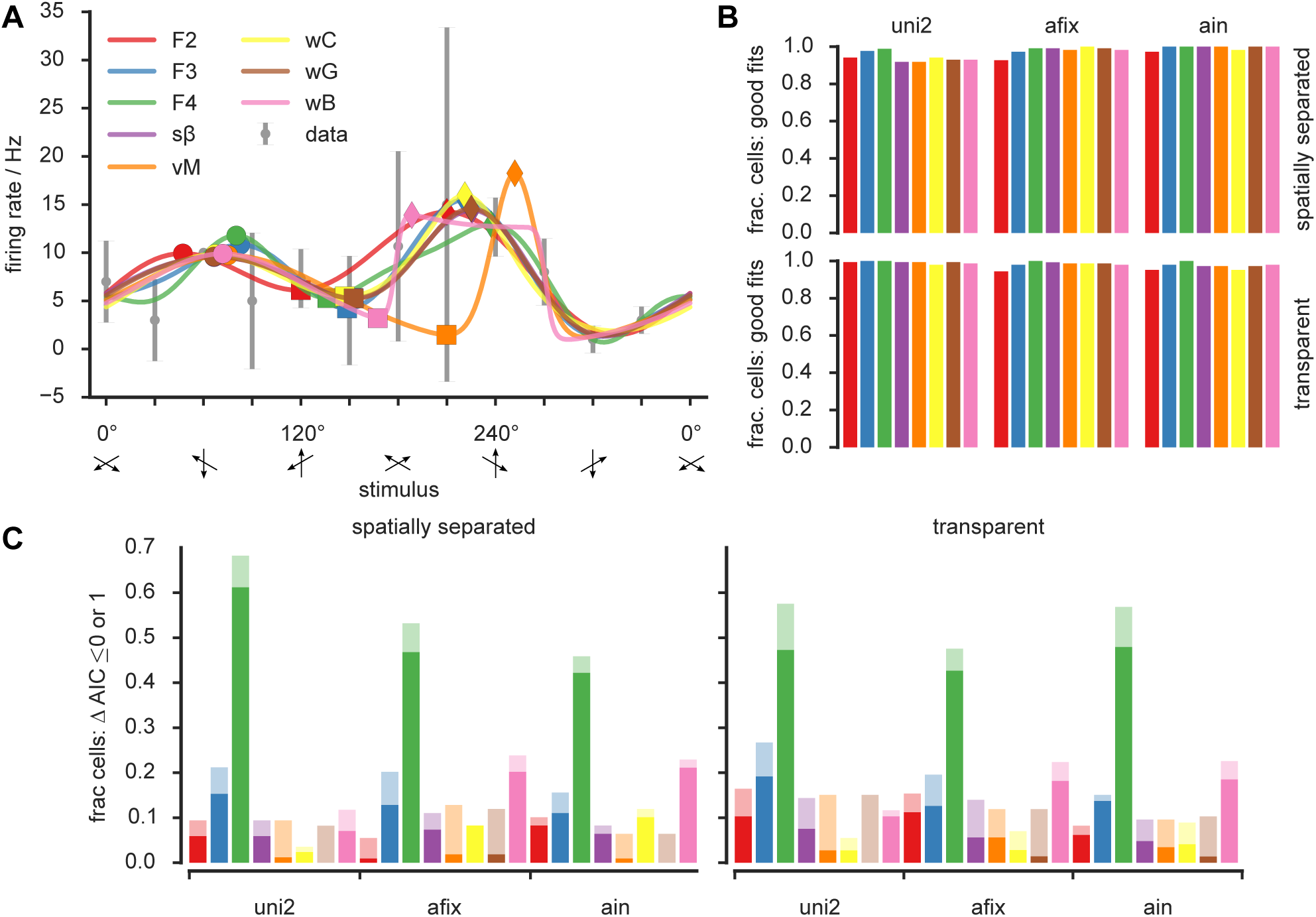
Many models are consistent with the data. **A)** Eight model functions were fitted to an example tuning curve (gray error bars denote std.) from the transparent afix condition. Due to the tuning curve’s large error bars all models provided good fits even though they clearly differ. **B)** According to a goodness-of-fit score (see main text) all eight models provided good fits for almost all cells, independent of experimental condition and paradigm. **C)** The Akaike Information Criterion AIC was thus employed to select the best (ΔAIC=0) or at least close to best (ΔAIC ≤) model for each cell. The fraction of cells for which each model constitutes the respective best or almost best model is illustrated with full and light bars. No model was chosen for all cells, still the most widely selected model was the fourth order Fourier series (*F* 4). Both of these facts mirror the high heterogeneity in the data that is hard to capture in a single tuning curve shape. **Color code** red: 2nd order Fourier (F2); blue: 3rd order Fourier (F3); green: 4th order Fourier (F4); violet: symmetric Beta (s*β*); orange - von Mises (vM); yellow: wrapped Cauchy (wC); brown: wrapped Gaussian (wG); pink: wrapped generalized bell-shaped membership function (wB).

We rigorously quantified goodness-of-fit by evaluating a score *Q*, giving the null hypothesis probability to observe by mere chance a sum of squared errors larger than in our fit. Small values of *Q* thus indicate poor fits (see Materials and Methods). We termed a fit “good” whenever *Q >* 0.1 but even lower values have been considered acceptable elsewhere [20]. We computed goodness-of-fit for every model type and for every cell, and we assessed the fraction of cells for which each given model type provided a fit deemed to be good. Fig. 3B shows that—at least according to the *Q* measure—nearly all cells could be well fitted by every model function, in every experimental condition. This statement still holds even when adopting more conservative thresholds for goodness-of-fit testing. As detailed in Supporting S1 Fig, even for threshold criteria as stringent as *Q >* 0.7, nearly all models provided good fits for more than 80% of the cells in most conditions.

Thus, for a majority of cells, goodness-of-fit alone was not enough to select the best model. We therefore adopted a model comparison approach and calculated the Akaike information criterion AIC [21] for different models (see Materials and Methods).

Smaller AIC values indicate better reproduction of the data by the model. Differences between AIC values for different models quantify how much information is lost describing the data with the model with a higher AIC value compared to the model with a smaller AIC value. Let, for a given cell, AIC_min_ be the minimum AIC value across all tested models. We then compute for each model *m* the relative information loss ΔAIC(*m*) = AIC(*m*) AIC_min_. Hence, for the best model ΔAIC = 0.

However—as a conventional rule of thumb—models with a small increase (ΔAIC *<* 1) should not be ruled out by model comparison but rather considered as equally good contenders for the “best fit” [22].

We show in Fig. 3C, the fraction of cells for which every tested model was evaluated as the “relative best”, i.e. obtained a value of ΔAIC = 0, as well as the fraction of cells in which it scored as an “equally good contender” with ΔAIC *<* 1. The outcome was qualitatively similar across all conditions. There was not a single model which scored systematically at the top for all cells, but each of the eight tested model types scored ΔAIC = 0 at least for a fraction of neurons. Even considering the softer criterion of ΔAIC *<* 1, none of the models appeared to be good enough to be used to fit all cells. Interestingly, for all experimental paradigms, Gaussian fits only quite rarely scored as the relative best (6–15 %, depending on paradigm and condition). On the contrary, Fourier series fits were the more frequent winners (72–100 %, taken together Fourier series of all used orders).

Some studies suggest that when the number of available samples is small, the corrected Akaike information criterion AICc [22] (see Materials and Methods) should be preferred to the AIC. This AICc penalizes models with a larger number of parameters more then the AIC already does (i.e. it implements a sharper “Occam’s razor”). We thus repeated the same analysis of Fig. 3C replacing the AIC with the AICc. Results are presented by the Supporting S2 FigA, which also presents the full statistical distributions of the observed AIC and AICc values (S2 FigB-C). It turned out that the best model according to AICc was always one with few parameters: either four or five in the uni condition and five in the afix and ain conditions. This indicates that models with only few parameters are enough to describe our highly irregular data. While no clear winner emerged in the unidirectional paradigm, the second order Fourier series fit model clearly outperformed the other models in the bidirectional paradigms, due to its reasonable fidelity in rendering the shapes of the measured tuning curves, combined with a smaller number of parameters.

In summary, model comparisons show that no single model can fit all cells equally well and that more than one model should be used when looking for the continuous interpolation of discretely sampled noisy tuning curves. Among parametric models tested (using both the AIC and the AICc criterion), Fourier series (rather than the commonly used Gaussian curves) tended to be the relative best in a larger number of cases. This reflects the substantial diversity of tuning curve shapes present in our representative dataset, since Fourier series do not have a single shape, but can faithfully render very dissimilar circularly wrapped profiles.

### Intermezzo: how to compare the shapes of different parametric fits

When fitting a model to data-points (**X**, **Y**) (such as a Gaussian profile **Y**(**X***, a, b, c, d*) = *a* exp(.5 (**X**–*c*)^2^*/b*^2^) + *d*), the set of the parameters of the model (*a, b, c, d* in this case) form a vector of descriptors of the shape of the relation linking **X** and **Y**. In the case of the Gaussian, indeed, the parameter *a* represents the peak amplitude, *b* the peak width, *c* the *x*-position of the peak amplitude and *d* the baseline level. For this reason, the shape of tuning curves and its alterations have often been analyzed in terms of the values of the parameters of a fitted model [3, 23]. However comparisons between fitted parameters are feasible only if a same underlying parametric model is used to fit tuning curves for all cells and in all experimental conditions. We have seen, on the contrary, that selecting a unique all-encompassing model to describe tuning might not be the best choice. Furthermore, fitted parameters do not have a direct geometric interpretation for every model. For instance, for Fourier series models, small changes of the internal parameters can lead to large changes in the resulting shape.

In order to compare between tuning curve shapes generated by different models in an intuitive way, we introduced generalized descriptive features that do not correspond to model parameters, but are extracted directly from the fitted curves through appropriate algorithmic rules (Fig. 4A).

**Figure 4.**
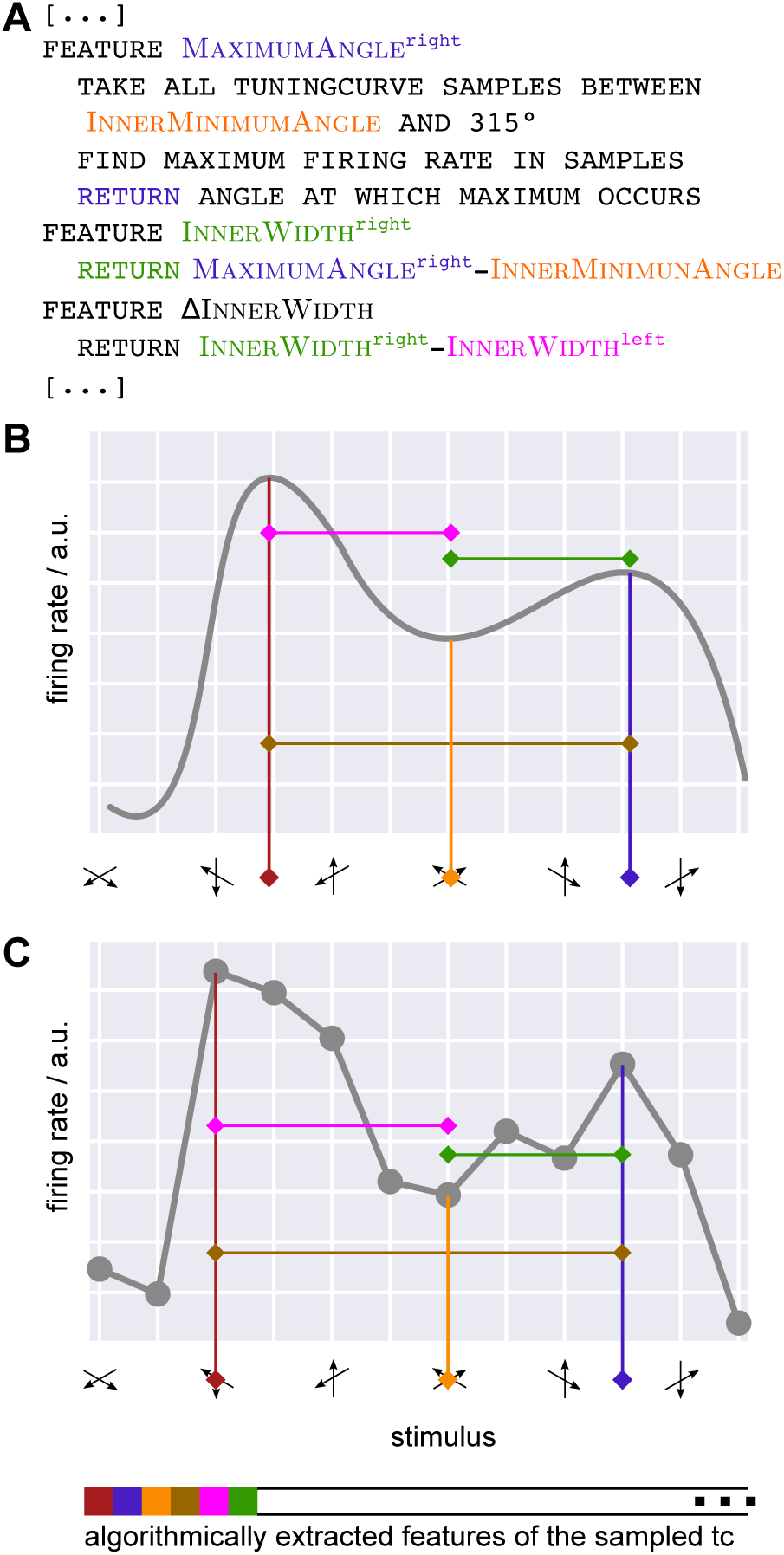
Alternative to fitting: algorithmic features. **A)** We describe all tuning curve properties of interest by algorithmically extracted features, as opposed to model parameters. The panel gives pseudocode for three selected features, illustrated also in panels B,C in corresponding colors. **B**) Feature extraction algorithms take as input only the sampled fitted tuning curve. Features are thus defined independent of a model function, allowing for their comparison between models. Furthermore, all aspects of tuning curves can be described by suitably chosen features such as Maximum^left^ (dark red), InnerMinimum (orange), Maximum^right^ (blue), (occuring at positions MaximumAngle^left^, InnerMinimumAngle and MaximumAngle^right^, respectively), InnerWidth^left^ (pink), InnerWidth^right^ (green). **C)** Due to their algorithmic nature feature extraction rules can equally well be applied directly to the coarse measured trial-averaged tuning curve. Thereby, tuning curve properties can be described and analyzed without referring to the fit.

The first step for the extraction of descriptive features is to select a certain number of points sampled along a fitted tuning curve profile and to note their coordinates (**X**, **Y**). Since the fitted profile is *continuous*, the number of selected points can be made arbitrarily large (unlike the number of actual measured data-points), but it is important to stress that the feature extraction approach that we introduce here always operates on a *discrete* set of points (a fact that will later allow us to apply it directly to the data-points themselves).

The second step is to apply the desired feature extraction rule to the sampled points. Consider, for example, a unimodal tuning curve, for which we could extract a feature GlobalMaximum, by finding the maximum *Y* coordinate among the points sampled along the profile. Analogously, we could introduce a feature MaximumAngle, corresponding to the preferred direction, by finding the *X* coordinate of the point whose *Y* coordinate corresponds to the feature GlobalMaximum. These and other features can easily be generalized to the case of bimodal tuning curves (cf. Fig. 4B). In this case the evaluation rules would be modified to limit the search just to the right (left) half of the sampled points to identify peak positions and amplitude of the right (left) peak respectively. Note that these features are based on points sampled along the fitted profiles, rather than on the *parameters* of these fitted profiles, which they reflect therefore only in a highly indirect manner.

Describing the extraction of a peak amplitude and position as a feature extraction procedure could be seen as a (generally non-linear) projection method [24] through which a continuous—or, at least, finely discretized—tuning curve shape is converted into a vector with a much smaller finite number of entries. This might seem unnecessarily complex, however there are several good reasons for doing so.

First, computed features may but do not need to mirror classic model parameters. For example, while the feature MaximumAngle would be equivalent to the parameter *c* in the case of a Gaussian fit, Fourier series don’t have any parameter directly reflecting peak position. More generally, all kind of convenient features can be designed, independent of the underlying model and tailored to describe specific shape aspects, such as, e.g., in a bimodal tuning curve, the position InnerMinimumAngle of the location of the lowest response between the two maxima —not necessarily centered between the two peaks— or the shape of the peak themselves whose OuterWidth and InnerWidth could differ, indicative of skewness or asymmetries. Table 1 gives a list of selected features, Supporting Tables S1-S4 provide the full list of features that we used as well as the detailed algorithm by which they were computed.

**Table 1.**
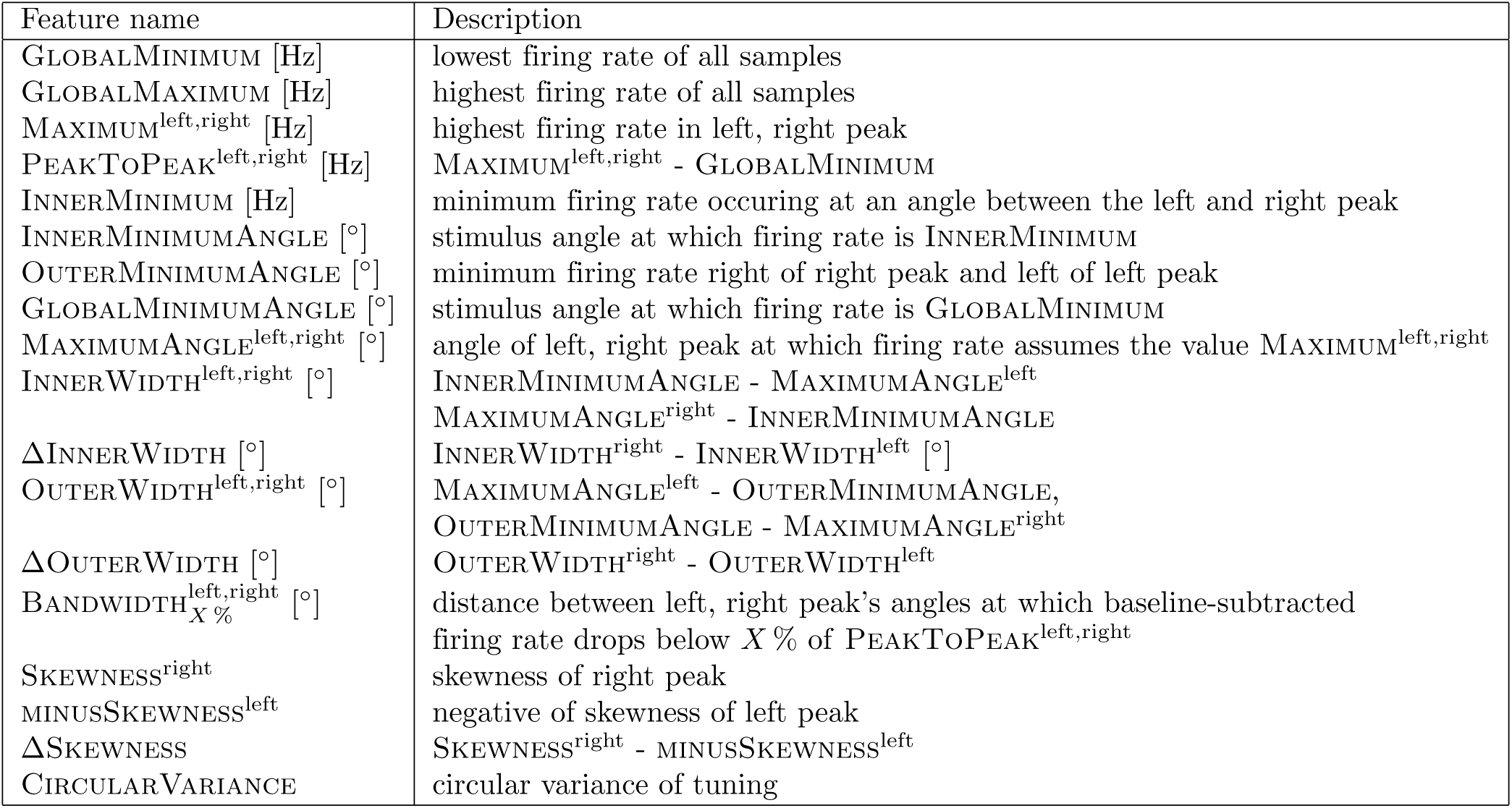
Selected features for the description of tuning curve shape. The complete list of features that we used as well as the algorithm for their computation can be found in Supporting Tables S1-S4

Second, as features are detached from the model itself, they allow a straightforward comparison of aspects of the tuning curves between parametrically incompatible models, which was a central aim in our study.

Third, as already anticipated, feature extraction rules can be applied directly—or with only minor modifications—to the observed data points themselves (Fig. 4C), without need of previously interpolating any continuous model curve. We will apply this direct approach, after completing our discussion of drawbacks inherent to model selection, by using feature extraction as a tool for comparing different fits.

### Different models can lead to different quantitative and qualitative results

As indicated above, Fig. 3A shows an example for which all eight model functions could be reasonably well fitted to the measured neuronal data. We parsed these eight fitted curves based on a common set of feature extraction rules (see Tables 1 and S1-S4). We then compared the extracted shape features between the models. In Fig. 3A the position of the right and left peak and the inter-peak minimum are highlighted, respectively, by circle, diamond and square symbols. Across the model fits the positions of the left peak differed by up to 35° (10 % of 360°) and the corresponding firing rates by up to 2 Hz (15 % of the maximum trial-averaged firing rate for the afix condition in this cell); the position of the inter-peak minimum varied by 90° (25 %), their firing rates by 5 Hz (31 %); the positions of the right peak by 64^*◦*^ (18 %), their firing rates by 5 Hz (35 %). The example neuron of Fig. 3A thus shows substantial differences between the shape features inferred when applying different models.

These differences at the level of a single cell generalized to a large fraction of cells in the dataset, creating significant differences between models at the population level. For each of the eight models we computed the distributions of the values of different features across cells. In addition we extracted features from a ninth model, denoted the best model (bM). In this bM case, we selected the best among the eight tested fits, as indicated by the ΔAIC = 0 criterion on a cell-by-cell basis, and for each cell we extracted features out of its specific best fit. Comparisons among the eight models and the ninth bM model are shown in Fig. 5A for nine selected features. In these nine matrices, a red entry indicates that the statistical comparison between the median values of the extracted features are significantly different between the two corresponding models (Kruskal-Wallis test, *p <* 0.05), while a blue entry denotes an agreement between the two models. Entries below and above the diagonal refer to feature comparisons for the spatially separated and the transparent paradigms, respectively.

**Figure 5.**
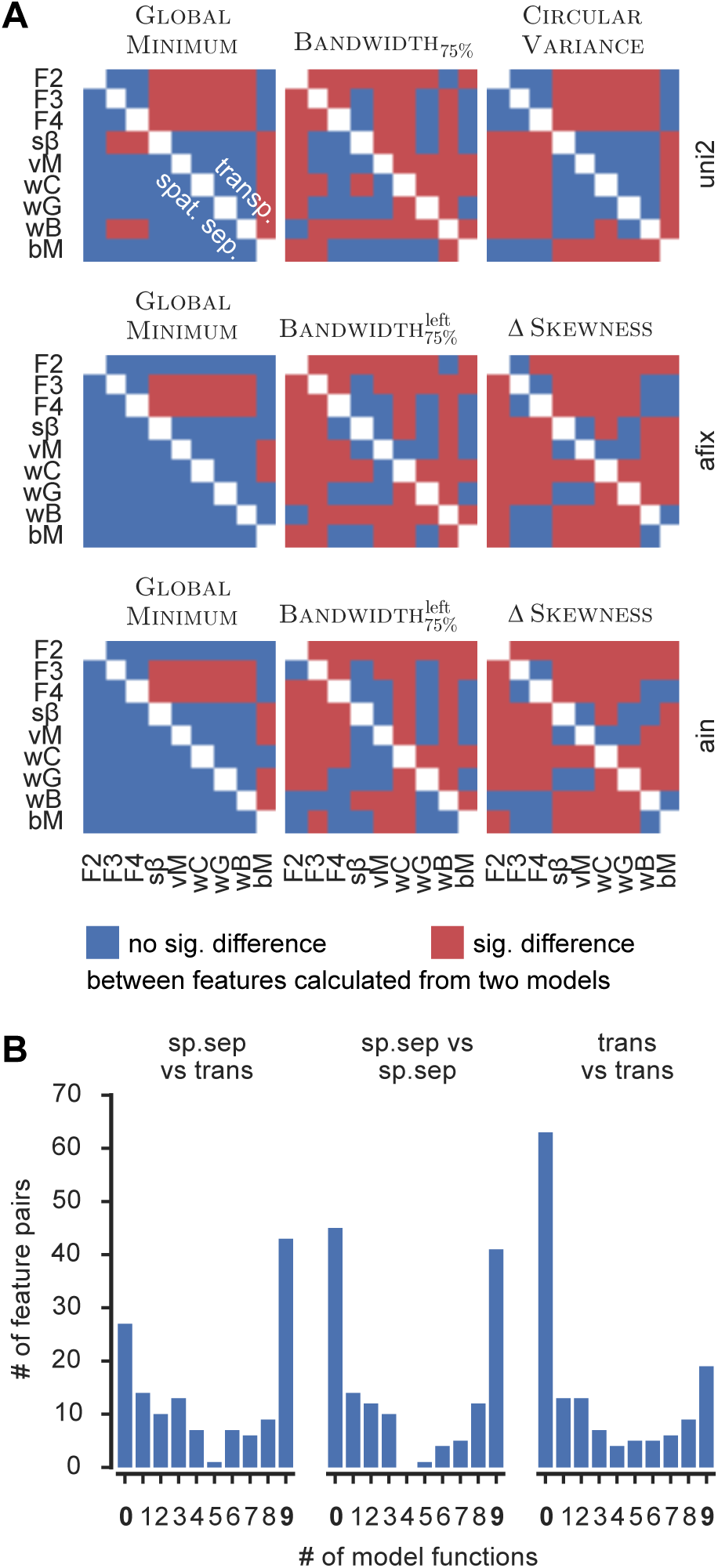
Effects found in the data depend on model. **A)** Nine features, measuring aspects of firing rate (first column), width (second column) and global shape (third column) in all three conditions were calculated for each cell on the basis of eight fitted models (model abbreviations are as in Fig. 3) and the “best model” (according to the ΔAIC=0 criterion, see main text; abbreviated “bM”). Red (blue) indicates a statistically significant (not sig.) difference between two models’ values of that feature. Results from the spatially separated and transparent paradigm are plotted below and above the diagonal, respectively. The panels indicate that, in general, models disagree on the value of a feature, and, in particular, might contradict the optimal (bM) model. Histograms count for all feature pairs (depending on their category “sp.sep vs trans”, “sp.sep vs sp.sep” or “trans vs trans”) the number of model functions that find a significant difference (“effect”) between the pair. While mostly all models agree (counts 0 and 9) there are also numerous cases in which the presence of an effect depends on the chosen model.

For some features, the median value did not change significantly for most model comparisons (for example the global minimum—GlobalMinimum—in all conditions of the spatially separated paradigm). For other features, there were marked differences between Fourier series and bM on the one hand, and the remaining models on the other hand (for example the circular variance, CircularVariance in the uni condition; but also the feature GlobalMinimum for the transparent paradigm). For yet other features, almost every model yielded different results (for example: the differences between the skewnesses, ΔSkewness, of the left and right peak in afix and ain; the 75 %-bandwidth of the left peak, 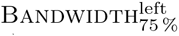, in afix; and the 75 %-bandwidth in the uni condition Bandwidth_75 %_). Importantly, in many cases there was a difference between bM and some of the other models.

Altogether, Fig. 5A demonstrates that the choice of model instead of another makes a difference, since it may lead to *quantitative* changes in the evaluated features.

However, it would be even more severe if these quantitative changes in the evaluation of specific features led to divergent *qualitative* conclusions on the comparison between experimental conditions. For instance, when studying attentional modulation effects, it is important to compare tuning curve peak amplitudes among, e.g. an afix and an ain condition. Say, for illustration, that we estimate in the afix condition, a median Maximum feature value of 30 Hz based on fits with the *i*-th model function, and a different median value of 40 Hz based on fits with the *j*-th model function. Let’s suppose that the median values for the *i*-th and *j*-th model in the ain condition read 42 and 45 Hz, respectively. Besides quantitative differences, it might happen that based on the *i*-th model we conclude that attention has led to a significant increase of the Maximum feature, but that this same comparison between the afix and ain conditions is not significant based on the *j*-th model. Thus, we would reach different conclusions on the effects of attention, depending on the chosen model. To check systematically for qualitative deviations in inter-condition comparisons, we performed comparisons between a large number of relevant feature pairs estimated from the nine different models (the eight tested model fit functions, supplemented by the bM model). We analyzed inter-model consistency for three different categories of comparisons: a feature from the spatially separate paradigm and the same feature from the transparent paradigm (this is possible for all defined features); a same feature taken from two conditions, e.g. uni vs afix, or afix vs ain (viable whenever the feature is defined for both conditions); or, two comparable features from a same condition (a list is given in supplementary table S5), e. g. Maximum^left^ vs Maximum^right^ for the peak firing rates of the two peaks of a bidirectional condition. For each tested feature pair we counted the number of fitted models for which the comparison was significant.

The histogram of these counts is reported, for different categories of feature pairs, in Fig. 5B. All histograms display a marked bimodal structure with two modes at the zero and nine counts values. These modes correspond, respectively, to the cases of complete agreement between models, i.e of a comparison which is never or always significant.

Since both these two cases were the most frequent, there was a robust tendency toward a qualitative agreement between the conclusions of different models. Crucially though, the gap between the two modes of these histograms was not empty, but there were frequent cases in which the significance of comparisons between two features in a pair depended on the adopted model. Thus, for all these feature pairs, the choice of a specific model for fitting tuning curves would have led to qualitatively divergent conclusions about the effects of attention. In particular, the reached conclusion might differ from the one drawn from the bM model, the one which was constructed as optimal on a cell-by-cell basis.

This makes it advisable to always fit tuning curves based on a bM mixture of models. However, the bM approach is particularly cumbersome to calculate. Furthermore, it is ill-defined. Indeed, given the high heterogeneity in the data, it is plausible that adding even more models to the list of candidates among which to perform the bM choice would lead to further qualitative differences. It seems therefore necessary to devise alternative strategies which completely avoid the questionable step of model selection itself.

### Feature extraction revisited: the direct method

Rather than relying on the extraction of tuning parameters from fitted data as illustrated above, rules for feature extraction can be generalized to operate on the experimental data points themselves. The main difference between a fitted profile and the empirical data points is a coarse angular resolution of experimental measurements that might potentially lead to a loss of precision of the extracted feature values.

However, this is a quantitative, not a qualitative difference, that does not prevent the application of the rule, as illustrated by Fig. 4C. We therefore extracted shape-describing features directly from the data for all cells in all experimental conditions and compared them with matching features estimated from different model-based fits.

Fig. 6 depicts the results of this comparison and Table 2 reports selected feature values obtained from the direct method and the corresponding values from the bM model (see Tables S6-S7 for a comprehensive list). We first focus on qualitative differences between the direct and the model-based approaches (Fig. 6A), before delving into quantitative differences (Fig. 6B). Fig. 6A follows an approach similar to Fig. 5B, however we now built distinct histograms for feature pairs which are significantly different based on the direct method (blue histogram) and feature pairs which are not significantly different based on the direct method (green histogram). As in Fig. 5B, we counted the number of fitted models with significant with feature differences. The blue (green) histograms—peaking at the maximum (minimum) model value count—indicate that when the direct method found a feature comparison to be significant (not significant), the most frequent case was that all nine (none of the) tested models also reached the same conclusion. Concomitantly, the left (right) tails of these histograms fell off quickly.

**Table 2.** Median values and significant differences for selected features. Numbers (numbers in parentheses) denote the median value over the population of cells from the corresponding paradigm and condition, calculated with the direct method (best model “bM”). Significant differences between conditions or paradigms (abbreviated here as “sp” and “tr”) are summarized (if present) in the line below the medians. <,, ≪ ⋘ denotes a p-value < 0.05, 0.01, 0.001, respectively, of a Kruskal-Wallis test applied to the feature values of the direct method. The condition listed before the relation sign had a smaller median than the one after it (if the medians were identical, means were compared). “NA” denotes features that were not defined for the corresponding condition.

| Feature | spatially separated |  |  | transparent |  |  |
| --- | --- | --- | --- | --- | --- | --- |
|  | uni | afix | ain | uni | afix | ain |
| GLOBALMINIMUM | 4.50 (4.14) | 7.14 (6.78) | 7.71 (8.09) | 1.96 (1.86) | 2.00 (1.53) | 1.93 (1.44) |
|  | sp uni < sp afix, sp uni <<< sp ain; tr uni << sp uni, tr afix <<< sp afix, tr ain <<< sp ain |  |  |  |  |  |
| GLOBALMAXIMUM | 31.00 (30.62) | 34.50 (35.15) | 38.50 (37.87) | 14.19 (15.01) | 14.00 (12.94) | 15.75 (15.93) |
|  | sp uni < sp ain; tr uni <<< sp uni, tr afix <<< sp afix, tr ain <<< sp ain |  |  |  |  |  |
| MAXIMUM<sup>left</sup> | NA (NA) | 29.50 (27.77) | 27.11 (27.89) | NA (NA) | 11.00 (10.37) | 12.22 (11.84) |
|  | tr afix <<< sp afix, tr ain <<< sp ain |  |  |  |  |  |
| MAXIMUM<sup>right</sup> | 31.00 (30.62) | 29.60 (30.34) | 36.29 (35.76) | 14.19 (15.01) | 12.00 (11.57) | 13.50 (12.80) |
|  | tr uni <<< sp uni, tr afix <<< sp afix, tr ain <<< sp ain |  |  |  |  |  |
| PEAKTOPEAK<sup>left</sup> | NA (NA) | 21.00 (20.89) | 17.14 (16.69) | NA (NA) | 9.08 (9.12) | 10.00 (10.06) |
|  | tr afix <<< sp afix, tr ain <<< sp ain |  |  |  |  |  |
| PEAKTOPEAK<sup>right</sup> | 25.50 (24.52) | 22.90 (23.58) | 26.60 (25.40) | 11.50 (11.09) | 9.50 (8.70) | 10.00 (9.96) |
|  | tr uni <<< sp uni, tr afix <<< sp afix, tr ain <<< sp ain |  |  |  |  |  |
| INNERMINIMUM | NA (NA) | 16.00 (15.29) | 18.80 (17.35) | NA (NA) | 4.00 (4.12) | 5.00 (3.68) |
|  | tr afix <<< sp afix, tr ain <<< sp ain |  |  |  |  |  |
| INNERMINIMUMANGLE | NA (NA) | 180.00 (174.00) | 150.00 (165.50) | NA (NA) | 180.00 (180.60) | 180.00 (188.35) |
| GLOBALMINIMUMANGLE | 60.00 (67.30) | 0.00 (12.80) | 0.00 (18.10) | 60.00 (52.10) | 30.00 (26.40) | 30.00 (21.10) |
|  | sp afix <<< sp uni, sp ain <<< sp uni; tr afix < tr uni, tr ain << tr uni; sp afix < tr afix |  |  |  |  |  |
| MAXIMUMANGLE<sup>left</sup> | NA (NA) | 120.00 (115.80) | 120.00 (120.10) | NA (NA) | 120.00 (113.70) | 120.00 (117.15) |
| MAXIMUMANGLE<sup>right</sup> | 240.00 (241.20) | 240.00 (240.20) | 240.00 (242.50) | 240.00 (244.35) | 240.00 (247.50) | 240.00 (247.10) |
|  | sp uni < sp ain; tr uni << tr ain; sp uni <<< tr uni |  |  |  |  |  |
| ΔINNERWIDTH | NA (NA) | 0.00 (8.70) | 30.00 (23.80) | NA (NA) | 0.00 (7.00) | 0.00 (3.45) |
|  | sp afix << sp ain; tr ain << sp ain |  |  |  |  |  |
| ΔOUTERWIDTH | NA (NA) | 30.00 (32.60) | 0.00 (21.30) | NA (NA) | 0.00 (2.10) | 0.00 (4.65) |
| BANDWIDTH<sup>left</sup><sub>75 %</sub> | NA (NA) | 90.00 (53.10) | 60.00 (50.50) | NA (NA) | 60.00 (47.90) | 60.00 (48.55) |
|  | sp ain < sp afix; |  |  |  |  |  |
| BANDWIDTH<sup>right</sup><sub>75 %</sub> | 90.00 (61.60) | 90.00 (54.80) | 90.00 (70.20) | 90.00 (60.90) | 60.00 (50.00) | 60.00 (50.40) |
|  | sp afix < sp uni, sp afix << sp ain; tr afix < tr uni;<br>tr uni < sp uni, tr afix < sp afix, tr ain <<< sp ain |  |  |  |  |  |
| ΔSKEWNESS | NA (NA) | 0.11 (-0.09) | -0.13 (-0.36) | NA (NA) | 0.58 (-0.10) | 0.42 (0.00) |
|  | sp ain < sp afix; sp afix << tr afix, sp ain <<< tr ain |  |  |  |  |  |
| CIRCULARVARIANCE | 0.46 (0.47) | NA (NA) | NA (NA) | 0.62 (0.59) | NA (NA) | NA (NA) |
|  | sp uni <<< tr uni |  |  |  |  |  |

**Figure 6.**
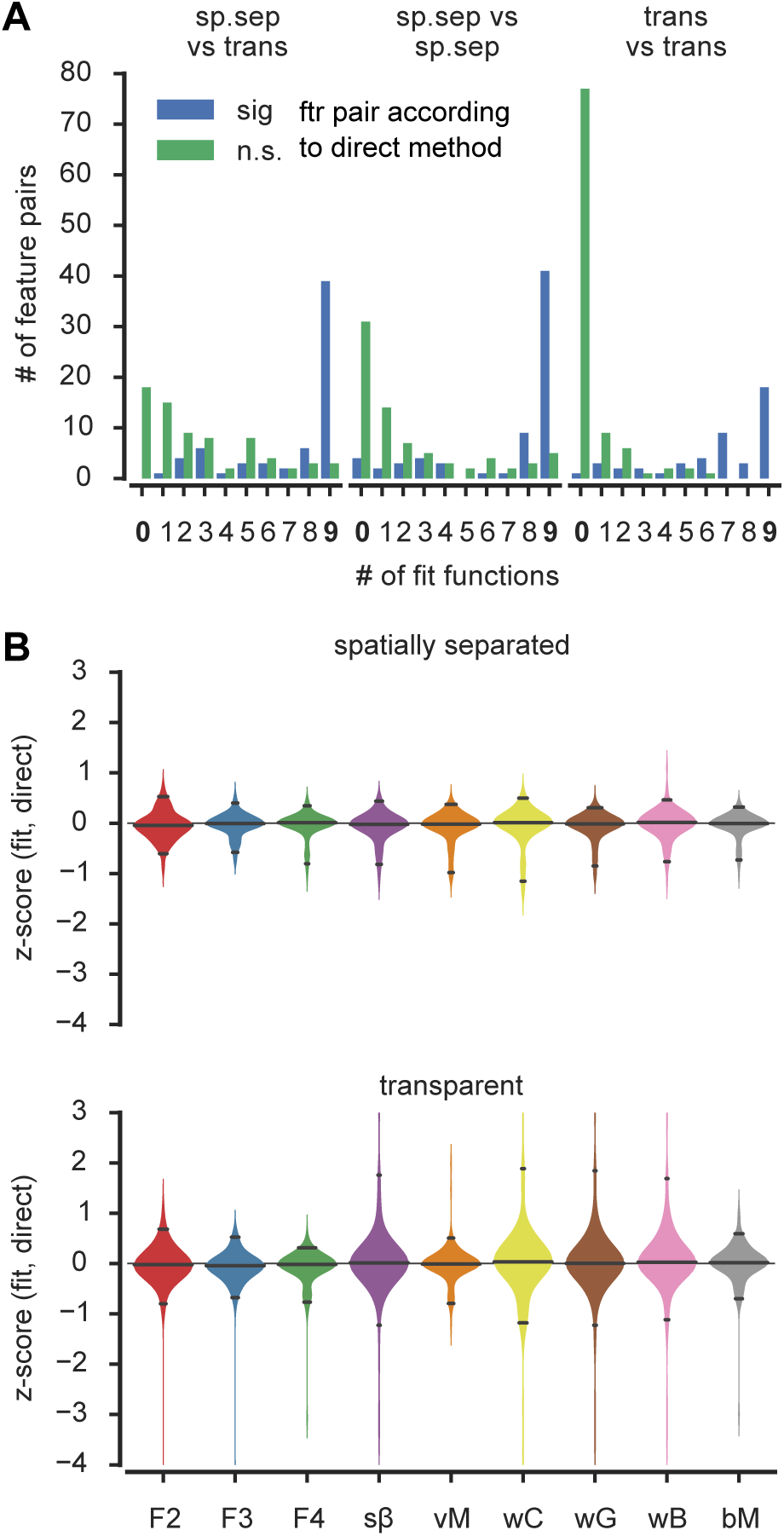
Direct method yields very similar results as fits. **A)** Layout is similar to figure 5B. Each bar from there was split into two, depending on if a considered feature pair was judged significantly different (blue) or not (green) when evaluated with the direct method. The panel illustrates a strong tendency to find a significant effect with either both the direct method and all nine models, or with neither the direct method and none of the models. **B)** *z*-scored quantitative differences between direct and fitted method’s feature values is less than one standard deviation for almost all features independent of the model indicating a considerable quantitative agreement between the methods. Solid lines in violines mark 2.5 %, 50 % and 97.5 % quantiles. Color code and model abbreviations are as in Figs. 3 and 5

Beyond the qualitative agreement, we also checked for quantitative agreements between features extracted by the direct and the model-based methods. We computed for each model *i* and for each feature *F* a *z*-score variable *z*_*i*_(*F*) = (mean(*F*)_*i*_ mean(*F*)_direct_) */* std(*F*)_direct_. The distributions of these *z*-scores over all the different features, for each different model function are plotted in Fig. 6B. For different experimental conditions and for all the model functions, the *z*-score distributions are centered on zero, indicating quantitative agreement between the direct and model-based methods. For the spatially separated paradigm all models gave results particularly close to the “direct” method (the difference was smaller than one standard deviation in almost all cases, i. e. *z* lay between -1 and 1, for at least 95 % of the extracted features). A relatively weaker quantitative agreement was observed for the transparent paradigm. For this paradigm, the Fourier series and the von Mises models gave the best agreement to the direct method. However, even in the case of the wrapped Cauchy model, which gave the worst agreement with the direct method, 88 % of the features had *z*-scores between -1 and 1.

In conclusion, we observed a qualitative and quantitative agreement between the direct method and the tested models. But note that the direct method makes less assumptions on the expected shape of tuning curves and does not conceal their heterogeneity.

### The direct method in action: Tuning curve modulations

After focusing on methodological aspects, we will now turn to the effects of different experimental paradigms, attentional conditions, and number of stimuli on the tuning.

We will concentrate on a narrow selection of significant feature variations (Kruskal-Wallist test with *p* < 0.05) revealed by the direct method, performing comparisons between conditions both within the same experimental paradigm and between different paradigms (Table 2 and Fig. 7). Supporting Tables S8-S9 provide then a complete list of significantly different feature pairs evaluated with the direct method and the best model.

**Figure 7.**
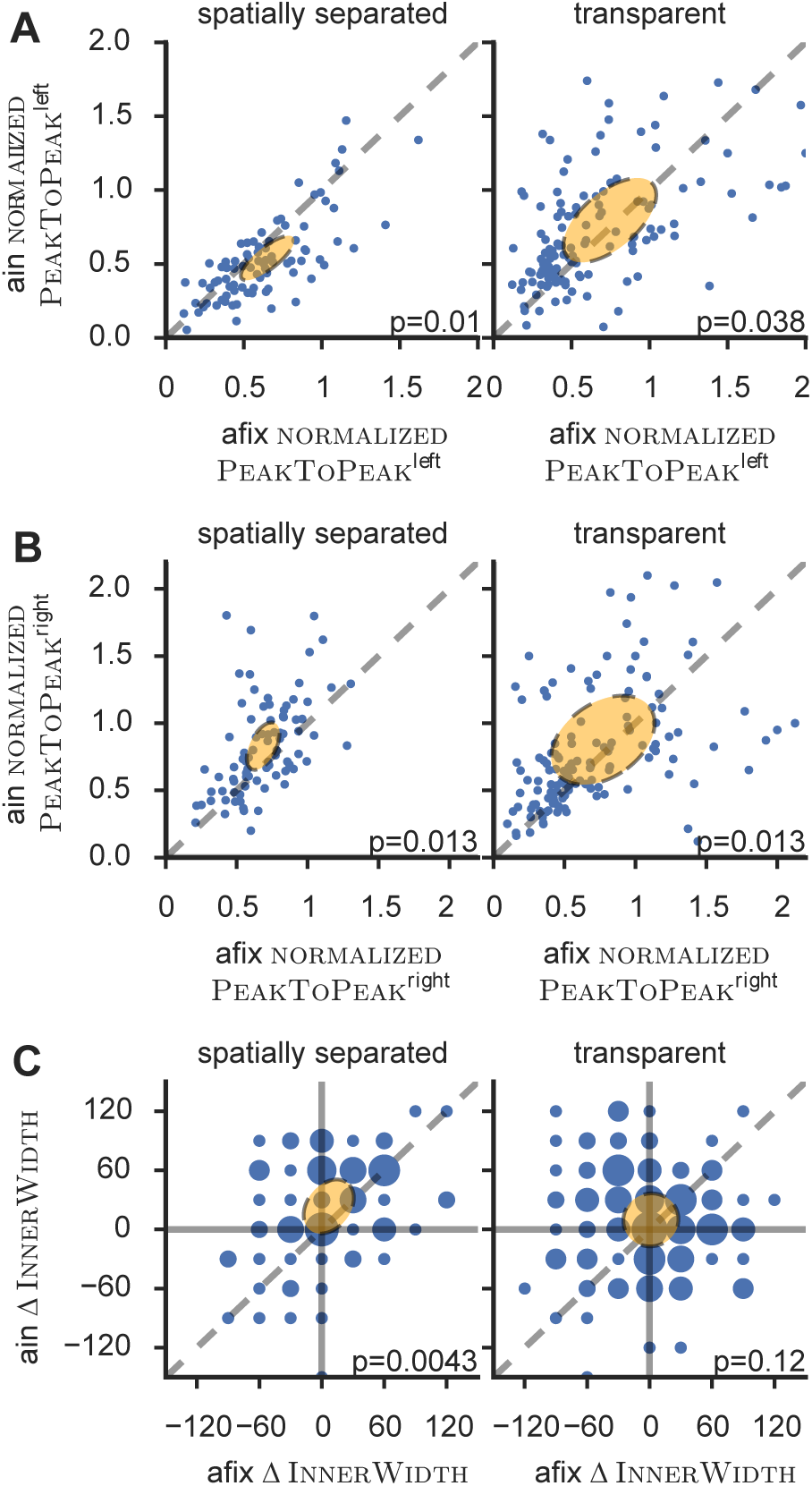
Effects of attention on tuning curves. Scatter plots of various features in the afix versus the ain condition. Orange error ellipses are centered on the mean feature values with half-axes corresponding to eigenvalues and -vectors of the feature-pair’s covariance matrix. Some outliers were omitted for better visualization. The indicated *p*-value in each panel corresponds to a Kruskal-Wallis test. **A)** Attention decreased (increased) the left peak—as measured by the feature normalizedPeakToPeak^left^—in the spatially separated (transparent) paradigm. **B)** Attention increased the right peak in both paradigms according to the feature normalizedPeakToPeak^right^. **C)** Attention significantly increased the difference between left and right peak’s inner width—ΔInnerWidth—only for the spatially separated paradigm. Size of circles in panel C illustrates density of points at each particular coordinate (note that values of ΔInnerWidth from the direct method are quantized in steps of 30^*◦*^due to the design of experimentally used stimuli). Altogether panels indicate that attention asymetrically expanded the right at the expense of the left peak for the spatially separated paradigm, but increased both peaks similarly for the transparent paradigm.

First, we evaluated tuning curves when one or two unattended stimuli were present in the receptive field, i. e. in the afix and uni conditions where attention was directed outside the receptive field (RF).

For the spatially separated paradigm, the peaks in the afix condition were smaller than in the uni condition. We monitored peak elevation over the baseline using the ad hoc engineered features PEAKTOPEAK^right^ and NORMALIZEDPEAKTOPEAK^right^ (analogous results hold for the left peak). These features quantify for each cell the variation between the right peak’s maximum and the response’s global minimum, which is normalized for the latter feature by the maximum firing rate in the uni condition (which is aligned, by convention, such that its peak overlaps the right peak in the afix condition). This NORMALIZEDPEAKTOPEAK^right^ feature decreased from 0.86 in the uni condition to 0.68 in the bidirectional stimulus afix condition (we report, here and in the following, sample median values). The same trend also held in the transparent condition, where NORMALIZEDPEAKTOPEAK^right^ decreased from 0.84 to 0.55, when superposing a second stimulus within the RF.

As detailed in the Methods section, in the spatially separate paradigm the uni condition was measured with attention directed to the fixation spot, whereas in the transparent case it was taken to be the cue-period of the ain condition, that is, attention was directed to the stimulus during the measurement. Accordingly, features regarding uni conditions cannot be compared between the two paradigms. Please note that this is due to the experimental design and does not limit the applicability of the method.

No significant differences were found between the amplitudes of the two peaks present in the afix condition, within both the spatially separated and the transparent paradigms, as monitored by the features MAXIMUM^left^ and MAXIMUM^right^.

We then evaluated the effects on tuning curves when deploying attention into the RF, i.e. in the ain condition.

We first monitored the emergence of amplitude differences between the two peaks of the tuning curve, computing, for instance, the feature ΔPEAKTOPEAK, i.e. the difference between PEAKTOPEAK^right^ and PEAKTOPEAK^left^. In the spatially separated paradigm, there was a significant increase of the amplitude difference between the attended and unattended peaks, with ΔPeakToPeak rising from 4 Hz in the afix up to 9 Hz in the ain condition, as a combined effect of a decrease of the left peak (NORMALIZEDPEAKTOPEAK changes from 0.6 to 0.5) and an increase of the right peak (from 0.68 to 0.76). In contrast, in the transparent paradigm, both peaks increased as an effect of attention (median NORMALIZEDPeakToPeak changed from 0.51 to 0.64 and from 0.55 to 0.70, for the left and right peaks, respectively). The different effect of attention on peak amplitudes in the spatially separate and transparent paradigms is also illustrated by scatter plots of the NORMALIZEDPeakToPeak in the ain vs the afix condition (Figs. 7A-B), where the cloud of points lies slightly below the diagonal for the unattended (left) peak in the spatially separated paradigm and slightly above it for the attended (right) peak in the spatially separated paradigm and for both peaks in the transparent paradigm.

Besides analyses of the amplitude and width of tuning curve peaks, the feature extraction approach allows the investigation of more general alterations in the response profile. The general shape of the attended peak differed between the spatially separate and the transparent paradigms. In particular the right peak was flatter (more platykurtic) in the spatially separate than in the transparent paradigm, as revealed by the median values of the feature 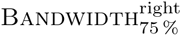 (see Table 1 for its definition), respectively, of 90^*◦*^ versus 60^*◦*^. A usually unreported effect of attention is illustrated in Fig. 7C, where we analyze variations of the InnerWidth features, i.e. the (absolute values of the) angular distance between each peak and the minimum between the peaks. Usually similar for both peaks in the afix condition, differences between the INNERWIDTH feature for the attended and the unattended peaks may signal interaction phenomena, such as, e.g., a tendency for the attended peak to re-absorb the unattended peak. For the spatially separated paradigm, the difference between the left and right INNERWIDTH, given by the compound feature ΔINNERWIDTH, increased significantly from 0° to 30°. On the contrary, no significant change was observed for the transparent paradigm. Note that the values of ΔINNERWIDTH are discretely quantized due to the coarse angular resolution of our measurements and the lack of interpolation in the direct method.

Conversely, the firing rate at the minimum between the peaks, monitored by the ad hoc feature normalizedInnerMinimum, increased significantly from 0.32 to 0.41, only for the transparent paradigm. For the spatially separated paradigm a trend in the same direction was also present, but was not significant.

Together these effects denote different shape alteration typologies for the two paradigms, which represent an asymmetric expansion of the attended at the expense of the unattended peak in the spatially separated paradigm and a symmetric, growth of both peaks for the transparent paradigm, increasing responses in the inter-peak dip.

In conclusion, the composition of multiple stimuli and the attentional state affected general global aspects of tuning curves, inducing characteristic and significant patterns of changes. The direct method allowed to isolate known effects of attention without need to resorting to any fit, and identified different patterns of attentional modulation for the two tested experimental paradigms. It also cast light on usually neglected aspects of tuning curve shapes such as peak asymmetries, which experimental condition and attention can also modulate, besides the most commonly studied effects on peak amplitude and width.

### Cell-and stimulus-specific aspects of attentional modulation

So far, the analyses have been based on responses averaged across all trials available for a given stimulus. While such an approach is very common, it is a simplification. Indeed, neuronal responses fluctuate strongly from trial to trial, which may be functionally relevant [25–28]. We therefore compared the distributions of responses across different attentional conditions, in a cell-by-cell and stimulus angle-by-stimulus angle fashion.

The cartoon in Fig. 8A illustrates this approach for a single cell. In the plot, each dots corresponds to the response of the cell in a given trial. The two experimental conditions (e.g., afix vs ain in the spatially separated paradigm) are represented in different colors. Comparing responses between any two conditions for matching stimuli, the large trial-to-trial variability stands out, as evident by scanning vertically the clouds of colored dots for any given fixed position on the horizontal axis (stimulus configurations). Accordingly, for some stimuli, the trial ensembles of responses may be significantly different between the two conditions (for example, in the cartoon of Fig. 8A, at the 60° stimulus), while for other stimulus configurations, the trial ensembles will not (for example at the 240° stimulus, in the cartoon).

**Figure 8.**
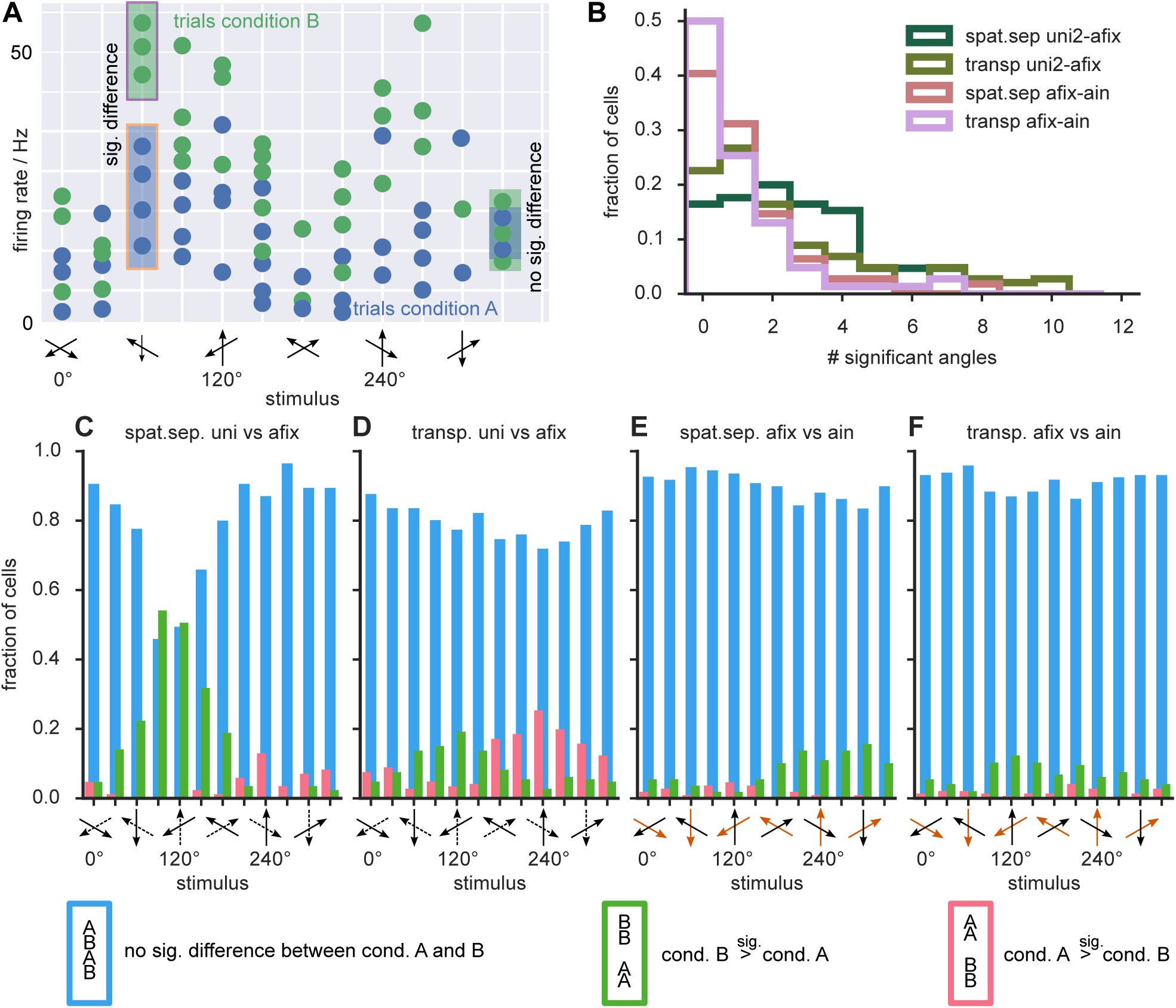
Effects of adding a second stimulus or attention to the receptive field. **A)** Each dot in this cartoon (not based on measured data) represents the observed spike count in one trial. For a given stimulus, spike count distributions can differ between experimental conditions either significantly (e. g. at 60°) or not (e. g. at 240°). **B)** Distribution of the proportion of cells with a significant difference between conditions for a given number of stimuli (maximum 12). The green histograms represent the two conditions where a second stimulus was added and pink histograms the conditions where attention was switched. **C-F)** Histograms show the stimulus-dependent fraction of cells with a non-significant response modulation (blue), a significant response enhancement (green) or response suppression (pink). The dotted and orange arrows along the *x*-axes in E and F indicate the RDP direction not present in the uni condition and the attended RDP in ain condition, respectively. Across the population a second stimulus tended to increase firing rates around 120°(C,D) and to decrease them around 240°. Attention asymmetrically affected the left and right peak in the spatially separated paradigm (E) whereas it symmetrically increased both peaks for the transparent paradigm (F). These stimulus-specific changes were compatible with the results of the direct method discussed in the text.

For each given cell we identified subsets of stimulus directions for which attention caused a significant response modulation (*p <* 0.05, two-sample Kolmogorov-Smirnov test). We found that these stimulus subsets were highly cell-specific, with different cells exhibiting robustly significant attentional modulations at different angles, not necessarily concentrated in proximity of a specific attended direction, but scattered for each cell over the entire range of possible stimuli (see Supporting Fig. S4 FigA). Correspondingly, we will refer to these stimulus-resolved significant differences, assessed at the single cell level as *specific effects*. While the statistical power of this analysis at the single cell level was limited by the small number of trials (see Supporting Fig. S4 FigB), the population level showed narrow stimulus ranges for which the fraction of significant specific effects were larger.

Despite the irregularity and large inter-cell variability of the significance patterns of specific effects, some weak overlap at the population level could still be identified, identifying narrow stimulus ranges for which the fraction of cells manifesting a significant specific effect were larger. Fig. 8B,C and D show how the response profile of a cell was altered when adding a second stimulus component within its RF, i.e. when going from a uni to an afix condition. The green histograms in Fig. 8B show the frequency distribution of the number of stimuli with significant changes, for the spatially separated and the transparent paradigm. In the spatially separated paradigm, 29% of cells showed significant changes for four or more stimulus directions and 16 % of the cells did not show significant specific effects at any angle. The corresponding numbers in the transparent paradigm are 25 % and 23 %, respectively. This means that, for around a fifth of all cells, the entire tuning curves for the uni and afix conditions where statistically indistinguishable.

The cells for which the addition of the second stimulus caused no significant response modulation tended to be poorly tuned already in the uni condition (see Supporting Figs. S3 FigA,B). On the other hand, some cells with equally poor tuning in the uni condition nevertheless displayed significant modulations of their firing rate when adding the second stimulus component.

Fig. 8B,E and F show cell-and stimulus-specific effects of attention, by comparing the afix and the ain conditions. The pink histograms in Fig 8B show the results of such a comparison for the spatially separated and the transparent paradigms. 7 % of the cells in the spatially separated paradigm and 10 % in the transparent paradigm showed significant specific effects of attentional modulation in four or more stimulus directions. The majority of cells in the spatially separated paradigm (60 %) and 50 % of the cells in the transparent paradigm showed a significant specific effect of attention for *at least* one stimulus. Note that most cells for both paradigms showed a clear tuning profile. Only 23 % (36 %) of the cells lacking any significant attentional modulation for the spatially-separated (transparent) paradigm were among the cells irresponsive to a second stimulus (Supporting S3 FigC,D).

Figs. 8C-F also show the stimulus directions for which significant specific effects of the addition of a stimulus component or of the allocation of attention were more frequently observed. The blue bars indicate the fraction of cells without a significant response modulation for a given direction, while the green and pink bars indicate significant increases and decreases of responses, respectively. Not surprisingly, the most frequent significant response enhancement occurred for the direction bins around 90° in the spatially separate uni vs afix comparison, corresponding to the cases where the preferred or a similar direction was added to a single stimulus moving about 120° away from the preferred direction.

For significant attentional modulations between the afix and ain conditions response increases were more frequent than response decreases at all stimulus directions, except 90° and 120° in the spatially separate paradigm (where response decreases were more frequent) and 60° in the transparent paradigm (where response increases and decreases were equally rare). In addition, significant attentional modulations, occurred mostly for stimulus angles between 180° and 330° (in the spatially separate paradigm) and 90° and 270° (in the transparent paradigm). These observations on stimulus-specific effects are compatible with the previous observation based on the direct feature extraction method, in that, for the transparent paradigm, both peaks are positively enhanced, while, for the spatially separate paradigm, only the attended peak is boosted, but the unattended one tends to be depressed. In this way the analysis of specific effects can shed light on the cell-level genesis of global shape changes of the average tuning curves. The trial ensemble comparison analyses presented in this section manifest how significant gain modulations at the level of a cell population may arise from the contribution of specific effects which are only rarely significant at the single cell level.

## Discussion

We showed that the commonly used approach to analyze tuning curve by fitting an idealized model function to the trial-averaged data may be more problematic than usually thought. Indeed, when adopting a model-based approach, there is a clear danger to reach model-specific conclusions (cf. [9]), which would not be confirmed by selecting different, equally viable models and which may be spurious. Here, going beyond model fitting and remaining within a purely data-driven framework, we extracted information about tuning and its modulations directly from the measured data points, through the application of rules for the extraction of suitable features. The high flexibility in feature design provided an antidote against over-constrained angles of view, which may be inherited by the adoption of narrow models.

Previous works already explored possible improvements on conventional least-squares fitting when dealing with noisy tuning curve data [4, 5] and a wide alternative of possible functional models to fit, not only Gaussians [1–4], but also typical circular statistics distributions [9, 10], as well as Fourier series [6, 9]. Even the most sophisticated techniques, however, are not immune to the drawbacks inherent to any procedure assuming a common underlying statistical model. On the contrary, as already pointed out long ago [8] and further confirmed by our analyses, the “best model” may vary from cell to cell, making the problem of its selection conceptually ill-posed.

Yet, fitting still remains a practical tool to inspect tuning behavior in data, abstracting, at least as a first step, from the variety of tuning curve shapes present in any dataset. Although the tested models all give rise to bell-shaped tuning profiles, they differ in the geometry of the bells’ flanks and these differences might be relevant for fine stimulus discrimination [16, 29]. Therefore, whenever fitting is used, one should carefully explore the entire *set* of candidate models, rather than of a single model, as a defense against excessive model bias. Results from our direct method itself could be included as well in the tested mix of analyses. A common set of features could then be extracted through a set of shared operational rules to systematically identify patterns of (or lack of) consistency between the diverse considered approaches. Comparisons between experimental conditions which are found to be significant only for a narrow subset of methods should then be looked at with suspicion, and confirmed by additional independent verifications.

The novelty of our data-driven approach is, however, more substantial than just providing yet another “model-less model”. First, if the correct model function cannot be certified with certainty, much of the seemingly high precision achieved by model-based interpolation may be just an illusion. Looking at the data in an agnostic and democratic manner, our data-driven methods could assess the statistical significance of attentional effects, strongly localized in both stimulus-and neuronal spaces. In particular they highlighted that only about 40 %-50 % of cells—similar to some previous reports of object-based attention in V1 [30, 31]—were significantly modulated attention, among them some virtually unresponsive cells which would often be discarded in conventional model-based studies. It remains an open question whether these specific effects are an artifact due to the limited availability of information (too sparse sampling of stimuli, limited number of available trials, etc.) or if they can be related to the fine-scale synaptic structure of top-down inputs. Indeed, at the local circuit level there is evidence for an extreme functional specificity of wiring [32, 33] and the frontal eye field, one of the assumed source areas of attention [34], might provide not more than a couple of synapses to excitatory (but not inhibitory) neurons in V4 [35]. In addition, models have shown that random and sparse recurrent network architectures are compatible with highly heterogeneous tuning curves [36, 37]. In the context of the present study, it is enough to stress that such fine-grained specific attentional effects would remain hidden to any approach based on the fitting of a stereotyped smooth model to cell responses. Adopting a model-free characterization of neuronal responses may thus well be necessary to relate advances in connectomics with cell-level modulations of functional activation.

Another potential application in which data-driven approaches could prove to be qualitatively superior to model-based approaches is the study of how attention affects complex population codes [38] of tuned responses. Indeed, by comparing trial ensembles of dozens of simultaneously recorded neurons previous studies already suggested that noise correlations were essential for the attentional performance enhancements [39] and that feature attention is coordinated across hemispheres whereas spatial attention correlates only local groups of neurons [40]. We, on the other hand, had only single cell recordings available, but they revealed a high degree of heterogeneity in tuning which may be functional, not merely reflecting noise, but carrying relevant information [26–29, 41–43]. In particular, such single-cell “weird” modulations may build up in a coordinated manner to give rise to population-level representations of the attended stimulus with a higher quality of encoding or with better and faster decodability properties [44, 45]. Until now only very few studies have addressed the recording of the tuned response of many cells simultaneously [39, 40, 46, 47] but the fast pace of growth of the number of simultaneously recorded neurons [48] will certainly call for more detailed characterizations of tuned responses, such as the ones that our methods begin to provide.

Conventional model fitting methodology is restricted to the analysis of a model’s parameters thereby potentially overlooking some features of the tuning with high discriminatory power. We have circumvented this problem in that we analyzed a set of features describing a wider range of aspects of the data. In the extreme case one could set up an all-encompassing feature library and programmatically mine for the most relevant ones. A possible drawback of massive feature libraries may be the feature selection analogue of over-fitting, i.e. the inevitability that some statistical comparison will appear to be spuriously significant just in virtue of multiple comparison issues. However, even this “data dredging” [49] is legitimate when used as an explorative technique for the generation of hypotheses to be verified by further studies on independently acquired data-sets. As a matter of fact, with twenty-first-century neuroscience entering an age of “big data” and large-scale cooperation [50], feature selection [24], in which features with optimized classification relevance are engineered in a (semi-)unsupervised manner, will increasingly become a method of choice for machine-augmented data-set parsing and knowledge discovery.

To conclude, although we are still far from understanding the intricate circuit mechanism through which attention influences information representation, routing and processing in the brain, we hope that our general methodology will assist the interpretation and inspire the design of future experiments necessary to advance this research endeavor.

## Methods

### Experimental procedures

All animal procedures of this study have been approved by the responsible regional government office (Niedersächsisches Landesamt für Verbraucherschutz und Lebensmittelsicherheit (LAVES)) under the permit numbers 33.42502/08-07.02 and 33.14.42502-04-064/07.

The animals were group-housed with other macaque monkeys in facilities of the German Primate Center in Goettingen, Germany in accordance with all applicable German and European regulations. The facility provides the animals with an enriched environment (incl. a multitude of toys and wooden structures), natural as well as artificial light, exceeding the size requirements of the European regulations, including access to outdoor space.

All invasive procedures were done under appropriate anesthesia and with appropriate analgesics. The German Primate Center has several veterinarians on staff that regularly monitor and examine the animals and consult on any procedures.

During the study the animals had unrestricted access to food and fluid, except on the days where data were collected or the animal was trained on the behavioral paradigm. On these days the animals were allowed unlimited access to fluid through their performance in the behavioral paradigm. Here the animals received fluid rewards for every correctly performed trial. Throughout the study the animals’ psychological and medical welfare was monitored by the veterinarians, the animal facility staff and the lab’s scientists, all specialized on working with non-human primates.

Three out of four animals were used in follow-up studies. One animal was euthanized at the end of the study. The decision was made in consultation with the attending veterinarian. Euthanasia was performed using an anaesthetic overdose (Sodium-Pentobarbital (i.v.)).

Single-unit action potentials were recorded extracellularly from extrastriate cortical area MT of four male rhesus monkeys (Macaca mulatta), using two sets of covert attention tasks. Two of the animals were performing the “spatially separated” paradigm, the other two the “transparent” paradigm. For the duration of every trial the monkeys were required to maintain their gaze on a fixation point in the middle of a computer monitor, placed at a viewing distance of 57 cm. While the animal maintained fixation, either one or two moving RDPs appeared in apertures in the receptive field (RF) of a given cell, as well as in the opposite hemifield outside the RF. In case of two RDPs the direction of the RDP in aperture 2 was always shifted clockwise from the direction of the other RDP by 120°. The direction of motion of the RDPs were varied in steps of 30° to obtain a tuning curve. The angular difference of 120° was selected as bi-directional tuning curves are expected to have two peaks in this case [16]. In the transparent paradigm the two RDPs were fully overlapping, crating just one aperture, covering most of the RF whereas in the spatially separate paradigm the two apertures were smaller and non-overlapping, but both still fully contained in the RF.

In the transparent condition there always existed just one aperture resulting in a single “uni” response profile. In the spatially separate paradigm, on the other hand, presenting the stimulus in either one of the two apertures gave rise to different responses, “uni1” and “uni2”, where the latter refers to the condition in which the stimulus appeared in the to-be-attended aperture (in the ain condition). If not noted otherwise, we always analyzed the “uni2” condition in the data from the spatially separate paradigm, and for simplicity also refer to it as just “uni”.

For the spatially separate unidirectional attend-fix condition (Fig. 1E) the monkeys were instructed to direct attention to the fixation spot, after a delay one RDP appeared in one of the two non-overlapping apertures and the monkey needed to detect a change of color of the fixation spot in order to receive a liquid reward. The spatially separate attend-fix condition (Fig. 1A) was similar, but RDPs were presented in both of the apertures. In the spatially separated attend-in condition (Fig. 1B) a RDP in one of the apertures was presented as a cue (of 500 or 600 msec duration), indicating to the monkey the location and the motion direction of a stimulus to be attended in the course of the trial. After a delay (800 ms) RDPs appeared in both apertures, and the monkey had to detect a transient change of motion velocity in the cued aperture at a random time point till maximally 2.5 s after the stimulus onset while ignoring possible changes in the other (distracting) RDPs.

The transparent attend-fix (Fig. 1C) and transparent attend-in (Fig. 1D) differed from the corresponding spatially separate conditions only in that the two apertures in which the RDPs were presented overlapped.

We generally analyzed data from the response period, which was defined as the time window 200-700 msec after onset of the stimulus in the RF. However, as no distinct unidirectional condition was recorded for the transparent paradigm, we used the cue period of the attend-in condition (50-500 msec after the cue onset) as a proxy for the uni condition in this case. That means that in the transparent uni condition attention was directed to the stimulus, whereas in the spatially separate uni condition attention was directed to the fixation spot. Accordingly, the uni conditions cannot be directly compared between the two paradigms.

We had 109 and 146 cells in the spatially separate and transparent paradigm, respectively. Uni conditions were recorded for 85 out of the 109 cells in the spatially separate paradigm. In 3 cells of the transparent paradigm the afix rates were not recorded and, therefore, our analysis disregarded the afix condition of those cells.

### Tuning data pre-processing

Data analysis was performed using custom-written software in Python (available on request). We did not perform spike-density estimation, but all analyses of tuning responses were based on raw firing rates, either averaged over trials (for model fitting and data-driven feature extraction) or estimated within each trial independently (for trial ensemble comparisons). Cells were included in the analysis only if at least two trials were available for every recorded condition. For some cells of the spatially separated paradigm no uni conditions were recorded. These cells were generally included in the analysis and exempted only in calculations concerning the uni condition.

All tuning curves were conventionally aligned, such that the maximum firing rate of the uni condition corresponded to the angular coordinate 240°. Whenever uni conditions had not been recorded (this was the case for some cells of the spatially separated paradigm) the angular position of the maximum firing rate of the right peak in the afix condition was defined to be 240°.

## Fitted models

To analyse tuning curves we fitted several model functions to the trial-averaged firing response data. Each fit to each specific cell was fully determined by the chosen parametric model and by a vector 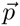 of model parameters. The wrapped Gaussian (*wG*) was given by:

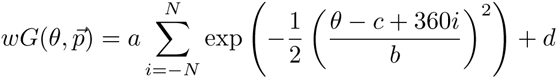

where 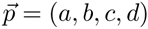 We always assumed *N* = 4 for wrapping.

The wrapped Cauchy (*wC*) function was given by:

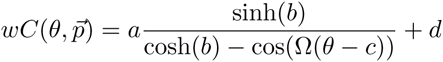

where 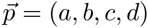and Ω = 2π/360.

The (modified) von Mises function (*vM*) was given by:

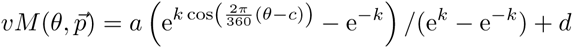

where 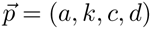

The symmetric Beta function (*sβ*) was given by:

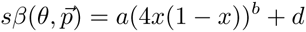

where *x* = (2*π/*360(*θc*) + π)/(2π) mod 1 and 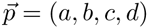

The wrapped generalized bell-shaped membership function (*wB*) was given by

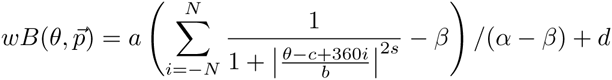

where 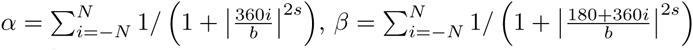, 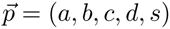 We always assumed *N* = 4 for wrapping.

All these functions are illustrated in Fig. 3A. In their basic form, they give rise to unimodal tuning profiles, as in the uni condition. To fit bimodal responses to composite stimuli, in the afix and ain conditions, we used a sum of two (identical) model functions, i.e.:

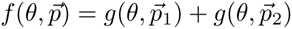

where *g* is either one of *wG, wC, sβ, wB* and the total parameter vector is 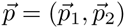 There was some redundancy between the parameter sets for the two peaks. For the first four functions *p*_1_ = (*a*_1_*, b*_1_*, c*_1_*, d/*2) and *p*_2_ = (*a*_2_*, b*_*c*_*, c*_2_*, d/*2), while, for the *wB* model, both 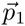 and 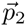 contained an additional component, *s*_1_ and *s*_2_, respectively. Hence, *f* had overall seven (*wG, wC, vM, sβ*) or nine (*wB*) free parameters.

In addition we also fitted Fourier series of order *n* = 2, 3, 4, given by:

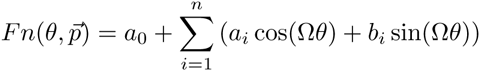

where Ω = 2π/360 and 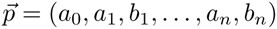 These Fourier series were fully determined by five, seven or nine parameters respectively, depending on their order *n* = 2, 3, 4. Fourier fits of unimodal or bimodal tuning curves shared a common functional form.

## Fitting methods

We used standard weighted non-linear least square fitting, relying on routines within Python’s SciPy package (http://www.scipy.org, sequential quadratic programming) for the minimization of the *χ*^2^ statistics. The applied initial conditions and boundaries therefore are listed in Supporting Table S10. An exception was given by Fourier series.

Let *p*_*i*_ be the *i*th component of the Fourier series’ parameter vector

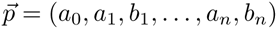 and *X*_*i*_(θ) be the *i*th compoment of (1, cos(Ω*θ*), sin(Ω*θ*)*,…,* cos(*n*Ω*θ*), sin(*n*Ω*θ*)). Then, an exact analytical solution to the least squares problem exists, which can be straightforwardly derived to be:

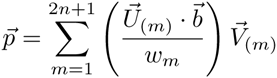

where 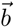 has components *b*_*i*_ = *y*_*i*_/*σ*_*i*_ 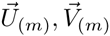denote the *m*th column of *U* and *V*, respectively. The matrices *U, V* and *w*_*m*_ form the singular value composition of matrix *A*, s.t. *A* = *U* diag(*w*_1_*,…, w*_2*n*+1_)*V* ^*T*^. The matrix *A*, in turn, has components *A*_*im*_ = *X*_*m*_(*θ*_*i*_)*/σ*_*i*_.

To quantify goodness-of-fit we also used a standard framework, as laid out in [20]. We assume that measurement errors in *y*_*i*_ are normally distributed. For model functions that are linear in their parameters —note, that in our library of models, this assumption holds only for the Fourier series *F n*—, the null hypothesis probability that the sum of squared errors is equal or larger than the observed *χ*^2^ is given by

*Q*(*K* − (2*n* + 1)/2, *χ*^2^/2=2) where 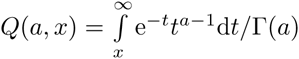 function and *K* is the number of independent samples (here, *K* = 12 tested stimulus ≤ directions). We use this quantity *Q* as measure for goodness of fit. If the probability *Q* is ≤ 10% we term the quality of the fit “bad”, otherwise we cannot rule out the hypothesis that the fit is an appropriate statistical models for our observations. For general non-linear models—for which the sum of squared errors cannot be expected to follow a conventional *χ*^2^ distribution—we evaluated approximately the goodness-of-fit *Q* statistics through a Montecarlo resampling approach (10000 replicas, cf. [20] for details).

## Model selection

We performed model selection based on the Akaike information criterion (AIC) [22, 51]. The information-theoretic quantity AIC gives the expected increase in uncertainty when using a certain model to describe the data rather than the “true” model. It can be computed from the sum of squared errors in the least-squares procedure according to

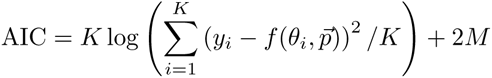

where *K* = 12 is once again the number of independent samples and *M* is the number of free parameters of the model function. Importantly, this formula is only valid in the limit of large *K*. Some studies [22, 51] therefore recommend to use a correction factor in the case in which 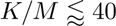. This corrected Akaike information criterion reads:

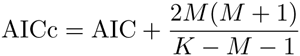

Such AICc converges to AIC for large *K* and mainly differs for it by applying a stronger penalty to models with larger number of free parameters.

## Features

Given a tuning curve *tc*(*θ*)—either as a discretized version of a fitted tuning profile, or directly by the vector of empirically observed average responses to stimuli with different directions—we characterized its shape in a non-parametric manner by calculating general *features*. A feature F is any map of the graph of the tuning curve *tc*(*θ*) onto some scalar number F : *tc↦* ℝ. Examples of features are the maximum of the tuning curve or the preferred direction (see Fig. 4 for an illustration). The complete list of features that we used is given in Supporting Tables S1-S4. Although we did not perform any feature clustering or redundancy elimination through e.g. factor analysis, willing in reality to maintain a library of features as wide as possible, we verified that most of the features bear complementary information, as hinted to by a sample distribution of pairwise correlations between feature values strongly peaked around zero (not shown).

Angular features were measured in degrees, ranging from 0^*◦*^ to 360^*◦*^ (with the exception of the feature GlobalMinimumAngle, for which we used the range −120^*◦*^ *≤* GlobalMinimumAngle ≤ 240^*◦*^) and coarsely quantized at just *K* = 12 equally spaced angular values. For each feature measured in units of Hz we also computed a normalized counterpart, denoted with the prefix normalized, by dividing the feature value by the global maximum firing rate found in the uni condition of the cell (if available).

We run statistical tests between *feature pairs* to search for effects of changes of experimental condition (transparent vs spatially separated, afix vs ain, etc.). For each of the two compared conditions we evaluated the values of the tested feature for each cell. We then performed two-way Kruskal-Wallis testing and dubbed a comparison as significant, whenever the *p*-value of this Kruskal-Wallis test was smaller than 0.05. All found significantly different feature pairs are listed in Supporting Tables S8 and S9, an excerpt in Table 2. The feature pairs reported herein are also the ones used in the systematic counting of significant comparisons reported in Figs. 5 and 6.

### Violin plots

Violin plots were calculated using Gaussian kernel density estimations with Scott’s rule (as implemented by www.scipy.org; [52]) for bandwidth estimation. Highlighted horizontal lines within the violin-shaped plot elements denote 2.5 %, 50 % and 97.5 % quantiles.

### Trial ensemble comparison

Beyond feature extraction we also compared directly vectors of firing rates measured across different trials for a same common cell and a same common stimulus. We then compared firing rate ensembles over trials for matching stimulus directions and cells across different experimental conditions, by means of a between-sample two-way Kolmogorov-Smirnov test. As for pairwise feature comparisons, we deemed a comparison between firing rate trial ensembles to be significant, whenever the *p*-value of this Kolmogorov-Smirnov test was smaller than 0.05. We call *specific effects* such stimulus-and cell-dependent effects of a significant change in condition revealed by trial ensemble comparison.

## Supporting Information

### S1 Fig

**The threshold value for** *Q* **above which to accept a fit is not critical**. Even for *Q*_thr_ > 0.7 more than 80 % of all cells lie above this threshold indicating a good fit.

### S2 Fig

**Model selection**. **A)** Layout as in Fig. 3C, but showing ΔAICc instead of ΔAIC. Also for this criterion, none of the models is always selected, although for afix and ain conditions second order Fourier (*F* 2) clearly performs best. **B,C)** Violinplots illustrating the distributions of ΔAIC (**B**) and ΔAICc (**C**).

### S3 Fig

**Cells not signficantly modulated by the addition of a second stimulus tended to be badly tuned but not vice versa whereas attentional modulation was unrelated to tuning.** We compared trial ensembles for a given stimulus between conditions. Dots mark tuning properties of cells with at least one (blue) and zero (red, brown) significantly different stimuli between **A,B)** uni and afix condition, and **C,D)** afix and ain condition. Red dots mark cells without any significant change in both comparisons. These cells (A,B) tended to be badly tuned, but there were also equally badly tuned cells sensitive to this manipulation. Directing attention to the receptive field (C,D) had no clear relation to tuning properties.

### S4 Fig

**Analysis of the statistical power for specific effects. A)** For each cell (*x*-axis) and stimulus (*y*-axis) colors indicate if there was a significant difference between the trials of the conditions marked in the title of each subplot. Colors are as in Fig. 8. **B)** We determined the impact of the number of trials available for our various conditions on the number of cells exhibiting significant changes between conditions. The plot shows how the number of cells exhibiting significant changes between conditions varied as a function of the number of trials included in the analysis (using the smaller of the two ensemble sizes for the *x*-value). The fraction of significant changes for all the tested condition changes showed a clear trend to increase with the number of included trials, possibly saturating when the number of trials reached about 8. Error bars denote standard-error of the mean. Note that points are only shown in this plot when we had a minimum of 10 samples, on average we had between 170 and 250 samples (depending on condition).

### S1 Table

**List of features defined for all tuning curves.** Each feature is calculated once for uni, once for afix and once for ain condition.

### S2 Table

**List of features defined only for uni condition.** Each feature is calculated only for uni condition.

### S3 Table

**List of features defined only for afix and ain condition.** Each features is calcualted once for afix and once for ain condition.

### S4 Table

**List of additional features comparing two conditions.** Conditions A,B can be either of uni, afix, ain.

### S5 Table

**Feature pair categories.** The features in each row were compaired against each other within one condition (uni, afic or ain) and for all conditions they are defined.

### S6 Table

**Spatially separated paradigm’s statistics for all features.** Table lists cell count, mean, standard deviation, minimum, 25 % quantile, median, 75 % quantle and maximum for all features when evaluated with the direct method (values from best model in parentheses).

### S7 Table

**Transparent paradigm’s statistics for all features.** Table lists cell count, mean, standard deviation, minimum, 25 % quantile, median, 75 % quantle and maximum for all features when evaluated with the direct method (values from best model in parentheses).

### S8 Table

**Significantly different feature pairs based on the direct method.** List of all significantly different feature pairs when evaluated with the direct method, as well as the corresponding *p*-value of the Kruskal-Wallis test, and medians, means and cell counts.

### S9 Table

**Significantly different feature pairs based on the best model.** List of all significantly different feature pairs when evaluated with the best model “bM”, as well as the corresponding *p*-value of the Kruskal-Wallis test, and medians, means and cell counts.

### S10 Table

**Initial conditions and bounds for least-squares-fits.** Model functions and their parameters are described in Methods subsection “Model functions”, number before model indicates if the model pertained to data from the one or two stimulus conditions. *u* = min_*i*_ *y*_*i*_ + 1.2 ptp*i y*_*i*_ where ptp*i y*_*i*_ = max_*i*_ *y*_*i*_min_*i*_ *y*_*i*_ and *y*_*i*_ is the set of all firing rates in the tuning curve. The values of 10.74, 2.41 and 100 for the width-parameter *k* of the s*β* model correspond to a half-width-at-half-maximum of 45°, 90° and 15^*◦*^. Likewise the values of *k* = 2.3, 0 and 20.34 for the vM model correspond to 45^*◦*^, 90° and 15°; As *k* = 0 would make the denominator in the definition of *vM* zero it was replaced by 0.001.

## References

1. Albright TD. Direction and Orientation Selectivity of Neurons in Visual Area MT of the Macaque. Journal of Neurophysiology. 1984 Dec;52(6):1106–1130. Available from: http://jn.physiology.org/content/52/6/1106.

2. Maldonado PE, Gray CM. Heterogeneity in local distributions of orientation-selective neurons in the cat primary visual cortex. Visual Neuroscience. 199613(03):509–516.

3. McAdams CJ, Maunsell JHR. Effects of Attention on Orientation-Tuning Functions of Single Neurons in Macaque Cortical Area V4. The Journal of Neuroscience. 1999 Jan;19(1):431–441. Available from: http://www.jneurosci.org/content/19/1/431.abstract.

4. Cronin B, Stevenson IH, Sur M, Körding KP. Hierarchical Bayesian Modeling and Markov Chain Monte Carlo Sampling for Tuning-Curve Analysis. Journal of Neurophysiology. 2010 Jan;103(1):591–602. Available from: http://jn.physiology.org/content/103/1/591.

5. Etzold A, Schwegler H, Eurich CW. Coding with noisy neurons: stability of tuning curve estimation strongly depends on the analysis method. Journal of Neuroscience Methods. 2004 Apr;134(2):109–119. Available from: http://www.sciencedirect.com/science/article/pii/S0165027003003935.

6. Wörgötter F, Eysel UT. Quantitative determination of orientational and directional components in the response of visual cortical cells to moving stimuli. Biological Cybernetics. 1987 Dec;57(6):349–355. Available from: http://link.springer.com/article/10.1007/BF00354980.

7. DeAngelis GC, Newsome WT. Organization of Disparity-Selective Neurons in Macaque Area MT. The Journal of Neuroscience. 1999 Feb;19(4):1398–1415. Available from: http://www.jneurosci.org/content/19/4/1398.

8. De Valois RL, William Yund E, Hepler N. The orientation and direction selectivity of cells in macaque visual cortex. Vision research. 198222(5):531–544. Available from: http://www.sciencedirect.com/science/article/pii/0042698982901122.

9. Swindale NV. Orientation tuning curves: empirical description and estimation of parameters. Biological Cybernetics. 1998 Jan;78(1):45–56. Available from: http://link.springer.com/article/10.1007/s004220050411.

10. Amirikian B, Georgopulos AP. Directional tuning profiles of motor cortical cells. Neuroscience Research. 2000 Jan;36(1):73–79. Available from: http://www.sciencedirect.com/science/article/pii/S0168010299001121.

11. Ringach DL, Shapley RM, Hawken MJ. Orientation Selectivity in Macaque V1: Diversity and Laminar Dependence. The Journal of Neuroscience. 2002 Jul;22(13):5639–5651. Available from: http://www.jneurosci.org/content/22/13/5639.

12. Dubner R, Zeki SM. Response properties and receptive fields of cells in an anatomically defined region of the superior temporal sulcus in the monkey. Brain Research. 1971 Dec;35(2):528–532. Available from: http://www.sciencedirect.com/science/article/pii/000689937190494X.

13. Born RT, Bradley DC. Structure and Function of Visual Area Mt. Annual Review of Neuroscience. 2005 Jul;28(1):157–189. Available from: http://www.annualreviews.org/doi/abs/10.1146/annurev.neuro.26.041002.131052.

14. Kozyrev V, Lochte A, Daliri M, Battaglia D, Treue S. attentional modulation of the tuning of neurons in macaque area MT to the direction of two spatially separated motion patterns. In: Society for Neuroscience. Chicago; 2009..

15. Lochte A, Stephan V, Kozyrev V, Witt A, Treue S. attentional modulation of the tuning of neurons in macaque area MT to the direction of transparent motion patterns. In: Society for Neuroscience. Chicago; 2009..

16. Treue S, Hol K, Rauber HJ. Seeing multiple directions of motion - physiology and psychophysics. Nat Neurosci. 2000 Mar;3(3):270–276. Available from: http://dx.doi.org/10.1038/72985.

17. Mardia KV, Jupp PE, editors. Directional Statistics. Wiley Series in Probability and Statistics. Hoboken, NJ, USA: John Wiley & Sons, Inc.; 1999. Available from: http://doi.wiley.com/10.1002/9780470316979.

18. Charalambides CA, Koutras MV, Balakrishnan N. Probability and Statistical Models with Applications. Auflage: 2003. ed. Boca Raton, FL: Chapman & Hall; 2000.

19. ϋbeyli ED. Adaptive Neuro-Fuzzy Inference Systems for Automatic Detection of Breast Cancer. Journal of Medical Systems. 2009 Oct;33(5):353–358. Available from: http://link.springer.com/article/10.1007/s10916-008-9197-x.

20. Press WH. Numerical recipes: the art of scientific computing. Cambridge [u.a.: cambridge Univ. Press; 2002.

21. Akaike H. A new look at the statistical model identification. Automatic Control, IEEE Transactions on. 197419(6):716–723. Available from: http://ieeexplore.ieee.org/xpls/abs_all.jsp?arnumber=1100705.

22. Burnham KP, Anderson DR. Model selection and multimodel inference: a practical information-theoretic approach. New York: springer; 2002.

23. Treue S, Trujillo JCM. Feature-based attention influences motion processing gain in macaque visual cortex. Nature. 1999 Jun;399(6736):575–579. Available from: http://dx.doi.org/10.1038/21176.

24. Guyon I, Elisseeff A. An introduction to variable and feature selection. The Journal of Machine Learning Research. 20033:1157–1182. Available from: http://dl.acm.org/citation.cfm?id=944968.

25. Arieli A, Sterkin A, Grinvald A, Aertsen AD. Dynamics of ongoing activity: explanation of the large variability in evoked cortical responses. Science. 1996273(5283):1868–1871. Available from: http://www.sciencemag.org/content/273/5283/1868.short.

26. Chelaru MI, Dragoi V. Efficient coding in heterogeneous neuronal populations. Proceedings of the National Academy of Sciences. 2008 Oct;105(42):16344–16349. Available from: http://www.pnas.org/content/105/42/16344.

27. Padmanabhan K, Urban NN. Intrinsic biophysical diversity decorrelates neuronal firing while increasing information content. Nature Neuroscience. 2010 Oct;13(10):1276–1282. Available from: http://www.nature.com/neuro/journal/v13/n10/full/nn.2630.html.

28. McDonnell MD, Ward LM. The benefits of noise in neural systems: bridging theory and experiment. Nature Reviews Neuroscience. 2011 Jul;12(7):415–426. Available from: http://www.nature.com/nrn/journal/v12/n7/abs/nrn3061.html.

29. Butts DA, Goldman MS. Tuning Curves, Neuronal Variability, and Sensory Coding. PLoS Biol. 2006 Mar;4(4):e92. Available from: http://dx.doi.org/10.1371/journal.pbio.0040092.

30. Roelfsema PR, Lamme VAF, Spekreijse H. Synchrony and covariation of firing rates in the primary visual cortex during contour grouping. Nature Neuroscience. 2004 Sep;7(9):982–991. Available from: http://www.nature.com/neuro/journal/v7/n9/abs/nn1304.html.

31. Poort J, Roelfsema PR. Noise Correlations Have Little Influence on the Coding of Selective Attention in Area V1. Cerebral Cortex. 2009 Mar;19(3):543–553. Available from: http://cercor.oxfordjournals.org/content/19/3/543.

32. Ko H, Hofer SB, Pichler B, Buchanan KA, Sjöström PJ, Mrsic-Flogel TD. Functional specificity of local synaptic connections in neocortical networks. Nature. 2011 May;473(7345):87–91. Available from: http://www.nature.com/nature/journal/v473/n7345/abs/nature09880.html.

33. DeBello WM, McBride TJ, Nichols GS, Pannoni KE, Sanculi D, Totten DJ. Input clustering and the microscale structure of local circuits. Frontiers in Neural Circuits. 20148:112. Available from: http://journal.frontiersin.org/journal/10.3389/fncir.2014.00112/full.

34. Moore T. The neurobiology of visual attention: finding sources. Current Opinion in Neurobiology. 2006 Apr;16(2):159–165. Available from: http://www.sciencedirect.com/science/article/pii/S0959438806000341.

35. Anderson JC, Kennedy H, Martin KAC. Pathways of Attention: Synaptic Relationships of Frontal Eye Field to V4, Lateral Intraparietal Cortex, and Area 46 in Macaque Monkey. The Journal of Neuroscience. 2011 Jul;31(30):10872–10881. Available from: http://www.jneurosci.org/content/31/30/10872.

36. Battaglia D, Hansel D. Synchronous Chaos and Broad Band Gamma Rhythm in a Minimal Multi-Layer Model of Primary Visual Cortex. PLoS Comput Biol. 2011 Oct;7(10):e1002176. Available from: http://dx.doi.org/10.1371/journal.pcbi.1002176.

37. Hansel D, Vreeswijk Cv. The Mechanism of Orientation Selectivity in Primary Visual Cortex without a Functional Map. The Journal of Neuroscience. 2012 Mar;32(12):4049–4064. Available from: http://www.jneurosci.org/content/32/12/4049.

38. Saproo S, Serences JT. Spatial Attention Improves the Quality of Population Codes in Human Visual Cortex. Journal of Neurophysiology. 2010 Aug;104(2):885–895. Available from: http://jn.physiology.org/content/104/2/885.

39. Cohen MR, Maunsell JHR. Attention improves performance primarily by reducing interneuronal correlations. Nature Neuroscience. 2009 Dec;12(12):1594–1600. Available from: http://www.nature.com/neuro/journal/v12/n12/abs/nn.2439.html.

40. Cohen M, Maunsell JR. Using Neuronal Populations to Study the Mechanisms Underlying Spatial and Feature Attention. Neuron. 2011 Jun;70(6):1192–1204. Available from: http://www.sciencedirect.com/science/article/pii/S089662731100434X.

41. Pouget A, Deneve S, Ducom JC, Latham PE. Narrow Versus Wide Tuning Curves: What’s Best for a Population Code? Neural Computation. 1999 Jan;11(1):85–90. Available from: http://dx.doi.org/10.1162/089976699300016818.

42. Seriès P, Latham PE, Pouget A. Tuning curve sharpening for orientation selectivity: coding efficiency and the impact of correlations. Nature Neuroscience. 2004 Oct;7(10):1129–1135. Available from: http://www.nature.com/neuro/journal/v7/n10/abs/nn1321.html.

43. Averbeck BB, Latham PE, Pouget A. Neural correlations, population coding and computation. Nature Reviews Neuroscience. 2006 May;7(5):358–366. Available from: http://www.nature.com/nrn/journal/v7/n5/full/nrn1888.html.

44. Tkacik G, Prentice JS, Balasubramanian V, Schneidman E. Optimal population coding by noisy spiking neurons. Proceedings of the National Academy of Sciences. 2010 Aug;107(32):14419–14424. Available from: http://www.pnas.org/content/107/32/14419.

45. Tkacik G, Marre O, Amodei D, Schneidman E, Bialek W, Berry MJ II. Searching for Collective Behavior in a Large Network of Sensory Neurons. PLoS Comput Biol. 2014 Jan;10(1):e1003408. Available from: http://dx.doi.org/10.1371/journal.pcbi.1003408.

46. Maynard EM, Hatsopoulos NG, Ojakangas CL, Acuna BD, Sanes JN, Normann RA, et al. Neuronal Interactions Improve Cortical Population Coding of Movement Direction. The Journal of Neuroscience. 1999 Sep;19(18):8083–8093. Available from: http://www.jneurosci.org/content/19/18/8083.

47. Stevenson IH, London BM, Oby ER, Sachs NA, Reimer J, Englitz B, et al. Functional Connectivity and Tuning Curves in Populations of Simultaneously Recorded Neurons. PLoS Comput Biol. 2012 Nov;8(11):e1002775. Available from: http://dx.doi.org/10.1371/journal.pcbi.1002775.

48. Stevenson IH, Kording KP. How advances in neural recording affect data analysis. Nature Neuroscience. 2011 Feb;14(2):139–142. Available from: http://www.nature.com/neuro/journal/v14/n2/abs/nn.2731.html.

49. Smith GD, Ebrahim S. Data dredging, bias, or confounding. BMJ : british Medical Journal. 2002 Dec;325(7378):1437–1438. Available from: http://www.ncbi.nlm.nih.gov/pmc/articles/PMC1124898/.

50. Sejnowski TJ, Churchland PS, Movshon JA. Putting big data to good use in neuroscience. Nature Neuroscience. 2014 Nov;17(11):1440–1441. Available from: http://www.nature.com/neuro/journal/v17/n11/abs/nn.3839.html.

51. Burnham KP, Anderson DR. Multimodel Inference: Understanding AIC and BIC in Model Selection. Sociological Methods & Research. 2004 Nov;33(2):261–304. Available from: http://smr.sagepub.com/cgi/doi/10.1177/0049124104268644.

52. Scott DW. Multivariate Density Estimation: Theory, Practice, and Visualization. Chicester: John Wiley & Sons; 1992.

